# Simulation of bacterial populations with SLiM

**DOI:** 10.1101/2020.09.28.316869

**Authors:** Jean Cury, Benjamin C. Haller, Guillaume Achaz, Flora Jay

## Abstract

Simulation of genomic data is a key tool in population genetics, yet, to date, there is no forward-in-time simulator of bacterial populations that is both computationally efficient and adaptable to a wide range of scenarios. Here we demonstrate how to simulate bacterial populations with SLiM, a forward-in-time simulator built for eukaryotes. SLiM has gained many users in recent years, due to its speed and power, and has extensive documentation showcasing various scenarios that it can simulate. This paper focuses on a simple demographic scenario, to explore unique aspects of modeling bacteria in SLiM’s scripting language. In addition, we illustrate the flexibility of SLiM by simulating the growth of bacteria on a Petri dish with antibiotic. To foster the development of bacterial simulations based upon this recipe, we explain the inner workings of its code. We also validate the simulator, by extensively testing the results of simulations against existing simulators, and against theoretical expectations for some summary statistics. This protocol, with the flexibility and power of SLiM, will enable the community to simulate bacterial populations efficiently under a wide range of evolutionary scenarios.

## 1 Introduction

Bacterial population genomics aims to reconstruct past evolutionary events and better understand the ongoing evolutionary dynamics operating in present-day populations. Demographic changes, selection, and migration are examples of processes whose genotypic signals remain in present-day populations. Trying to recover these signals from ever-growing sequencing data is a major goal of population genomics. In the context of epidemiological surveillance the inference of these types of events can be useful, since pathogens are known to undergo frequent demographic changes [1]. Similarly, a better understanding of the evolutionary forces operating in pathogen populations can help to inform public health policy [2]. For instance, one can assess the impact of a vaccination campaign or the efficacy of a new antibiotic on a given pathogen population [3], [4]. Beyond these clinical settings, bacterial population genomics can also be useful to describe natural population diversity [5].

Simulations are essential to population genetics [6]. They are useful for testing and validating population genetics methods (whether based on simulations or not), since they provide data generated by known evolutionary forces (unlike, typically, empirical sequence data). Notably, they can be used to assess the performance of statistical methods when assumptions are violated [7], [8]. They are also helpful for predicting the impact of an environmental change on a population, or the expected response to intervention [9], [10].

Many methods inferring past evolutionary events also rely on simulated data. In Approximate Bayesian Computation (ABC), probably the most famous likelihood-free inference framework in our field, simulations enable estimation of the posterior distribution of parameters of interest [11]. Other methods, based on machine learning, also require simulations to train a model to learn the mapping between input sequence data and evolutionary processes [12]. Increasingly, machine learning methods involve “deep learning” algorithms that hold great promise but require a large volume of simulated data [13]-[17]. Despite these many applications of simulations in population genetics, there are very few bacterial population genetics simulators, and those that exist do not cover many possible scenarios. In particular, existing bacterial simulators are coalescent-based simulators (`msPro` [18], `SimBac` [19], `FastSimBac` [20]), which means they are very fast and memory-efficient, but can model only a narrow range of evolutionary dynamics. For instance, these simulators do not allow selection, and in the case of SimBac, demographic changes cannot be simulated either. Simulation of complex selective forces together with demographic processes remains a difficult problem for coalescence-based simulators [19]; other coalescent-based simulators that are not specific to bacteria (e.g., `ms` [21], `msprime` [22]) suffer from the same constraints, and additionally most cannot simulate bacterial recombination (similar to gene conversion). On the other hand forward simulators such as SFS_CODE [23] enable complex models including varying demography and multiple types of selection; however, this software suffers from poor performance [24]. Yet computational efficiency is crucial for supervised methods trained on large simulated datasets, such as ABC, machine learning, and deep learning. Using forward simulation instead of coalescent-based methods can therefore be problematic, given that forward simulation has traditionally been several orders of magnitude slower than the coalescent. CoreSimul, a forward-in-time bacterial simulator with bacterial recombination and selection, was recently published [25]; however, it is designed for a different problem space than we are interested in here (simulation of different models of molecular evolution on a fixed genealogical tree of sampled individuals).

Here we present a method for simulating bacterial populations in a flexible and fast way, using a forward simulator called SLiM [26]. The SLiM forward simulation framework is becoming quite widely used due to its speed, power, and flexibility [27]-[29]. SLiM includes a scriptable interface with its own language, Eidos, which allows simulation of a wide range of evolutionary dynamics. The detailed instruction manual, combined with its helpful graphical user interface and its versatility, enable users to build simulation models tailored to their research. Simulation of bacterial populations, and haploids in general, is not supported intrinsically by SLiM, because every individual has two chromosomes. But because of its scriptability, it is possible to extend SLiM into this area. In this protocol, we will show the key techniques necessary to perform bacterial simulations. Following the SLiM manual’s convention, we will introduce the model implementation step by step together with the related concepts. Then we will show that the simulator behaves correctly according to the expected values of certain summary statistics under the Wright-Fisher model or by validating against other simulators, and that the model’s performance is good enough to allow numerous simulations to be run in a reasonable amount of time and memory. Finally, we will showcase a more complex model, based upon our method, in which we simulate bacteria growing on a Petri dish. This model is spatially explicit, representing the bacteria actually colonizing the dish, half of which contains an antibiotic that decreases their survival rate. Resistance mutations may emerge, substantially increasing the fitness of bacteria growing in the presence of antibiotic, while slightly decreasing fitness otherwise. This model illustrates the open-ended flexibility of SLiM.

We believe this work will open new avenues in bacterial population genetics by allowing researchers to go beyond the limitations of the coalescent, broadening the potential applications of simulation to a much wider range of evolutionary dynamics.

## 2 Methods, simulator and data

The bacterial simulator proposed here is based on SLiM, a powerful and efficient forward genetic simulator [30]. Thanks to its flexible scripting interface using the Eidos language, we were able to adapt SLiM to the simulation of bacterial populations.

SLiM provides two types of simulations: Wright-Fisher (WF) models, and models that go beyond the Wright-Fisher framework (non-Wright-Fisher or nonWF models). The Wright-Fisher model is based on many simplifying assumptions that are often not compatible with realistic scenarios such as structured populations, overlapping generations, etc. [26]. However, it is mathematically simple, allowing expectations for certain quantities to be estimated. This is particularly useful to validate the created simulator against the expectations under this model. The nonWF framework, on the other hand, is more individual-based, emergent, and realistic. It allows a greater breadth of possible scenarios to be simulated, but we cannot derive expectations of the same quantities. Thus, we will provide results for the same scenario under both models to confirm that they behave similarly (according to the WF expectations).

In the main text we will present the protocol for simulating bacterial populations with the nonWF framework, since it is a more powerful framework on which other users can build more complex scenarios. The underlying simulation, however, corresponds to a Wright-Fisher population, so we can compare to the theoretical expectations. The corresponding annotated WF script is available in a public repository (https://github.com/jeanrjc/BacterialSlimulations), along with the nonWF script detailed below. The different simulators used in this article are summarized in the table 1 below. To highlight the modeling steps that are specific to bacterial populations, we kept the underlying population history simple, with a single constant-size population and no selection, but those assumptions are trivial to relax in SLiM.

**Table 1:** Summary table of the simulators used in this paper. Phases 1 and 2 represent simulation steps in the order that they are performed. The third column gives brief comments regarding each simulator, while the last column gives an example of the degree of scenario complexity that can be simulated with each.

|  | Phase 1 | Phase 2 | Comments | Example |
| --- | --- | --- | --- | --- |
| ms / msprime / FastSimBac | Coalescent simulation | None | Ideal for simple simulations of basic scenarios | Migration between populations with demographic changes |
| SLiM with Wright-Fisher (WF) framework | Burn-in with ms | Forward simulation with SLiM, starting from burn-in output | Ease of implementation of slightly complex scenarios; fitness affects fecundity | Demographic changes and background selection |
| SLiM with nonWF framework | Forward simulation with SLiM and tree sequence recording | Recapitulation of the tree sequence using msprime | Wide range of possible scenarios; fitness affects mortality; faster and more accurate burn-in | Spatial model with environment-associated fitness for certain mutations |

### 2.1 Key concepts and definitions

#### 2.1.1 Horizontal gene transfer, recombination, and circularity

In bacteria, pieces of DNA can be exchanged between different organisms in a process called horizontal gene transfer [31]. When received, such a DNAfragment can be inserted in the host chromosome with the help of integrases, at a specific site if the fragment is not homologous to an existing chromosomal region. Alternatively, if the incoming DNA fragment *is* homologous, it will integrate into the host chromosome by a mechanism similar to gene conversion in eukaryotes [32]. This latter process is the bacterial recombination mechanism that we want to implement. We will use the term “gene conversion” to refer to gene conversion specifically in eukaryotes, and the term “bacterial recombination” to refer to the mechanistically similar process in bacteria. Note that bacterial recombination differs from simple recombination in eukaryotes (often called cross-over), in that mutations are not *exchanged* between two fragments of DNA; instead, the mutations are *copied* from one fragment to the other. Some coalescent simulators do implement gene conversion or bacterial recombination, such as `ms` [21] or `FastSimBac` [20], but they are otherwise quite limited, as discussed above. Our implementation of bacterial recombination also provides a slightly closer fit with reality, since we model the bacterial chromosome as circular. It has been shown that circularity can lead to different patterns, such as linkage disequilibrium decaying faster in linear genomes than in circular genomes [5], [33]. Although circularity is likely not important for the metrics and parameters shown in this study, including it is one less incorrect assumption when modeling bacteria.

#### 2.1.2 Burn-in

It is often desirable to start a simulation with a population which is at mutation-drift equilibrium. After 5×Ne generations in our simple scenario, the heterozygosity has reached more than 99% of the heterozygosity expected under mutation-drift equilibrium (see Annexes for demonstration). In forward simulations, the time spent to reach this equilibrium (5×Ne) is called “burn-in”. Because the effective size of bacterial populations is usually large, conduct-ingthis burn-in with a forward simulator would often require much more time than simulating the actual time period of interest, and this problem can make forward simulation of bacteria difficult or even infeasible.

To solve this issue, faster backward-in-time simulators can be used to simulate a population at equilibrium that serves to initialize the forward simulation. The nonWF model allows an elegant and efficient approach to this: we can combine SLiM’s tree-sequence recording feature with the recapitation feature of `msprime` [26] to manage burn-in. With this strategy, we can begin with forward simulation in SLiM, leaving the burn-in for later. At the end of the forward simulation, there is often no single common ancestor for the population; in other words, the ancestry tree of the underlying population has not yet coalesced. Recapitation will then simulate, backward in time, the addition of ancestral branches to produce coalescence, providing the needed burn-in ancestry after the fact. However, because `msprime` does not yet implement gene conversion, we cannot use bacterial recombination during burn-in for our nonWF model.

The WF model requires a different approach, because tree-sequence recording cannot be used; in the WF model SLiM cannot record HGT events in the tree sequence. In this case, we therefore have to simulate the entire population backward in time with `ms`, and load the generated diversity into SLiM to initialize its forward simulation. Because we simulate the entire population, it is not possible to use gene conversion at a significant rate, otherwise ms crashes; thus there is no bacterial recombination during burn-in for our WF model, either. As of now, for long simulations, it is thus not possible to have bacterial recombination during a coalescent-based burn-in; we analyse the impact of this limitation on simulations in the results section. For small populations, however, the burn-in can be simulated directly in SLiM. Finally, in certain situations a burn-in is not desirable (as in our Petri dish model).

#### 2.1.3 Simulation rescaling

Forward simulators remain computationally intensive, and bacterial populations can be very large. The effective population size of most bacterial species is on the order of 10^8^ – 10^9^ [34]. Depending on the task one wants to address, many thousands or even millions of simulations may be required. One way to reduce the computation time is to parallelize the simulations on a cluster, but it can remain costly. Another way is to rescale the model parameters such that *θ* = 2 × *Ne × μ* and related quantities remain constant [35]. For instance, we can decrease the size of the population by a factor of 10 while increasing the mutation and recombination rates by the same factor. The choice of the rescaling factor is at the discretion of the user, but one should keep in mind that excessive rescaling might lead to spurious results [35]. For instance, rescaling increases the rate of double mutation at a site, although it should remain rare [36]. Also, if the simulation involves a bottleneck, the user should make sure that the number of individuals remaining in the population after the bottleneck is not so small as to cause artifacts. A model of a bottleneck that reduces a population of 1000 individuals to 100 would lead to very different results if we were to rescale the model down to only 10 individuals before the bottleneck and one individual after! The rescaling factor must also be applied to the duration of the simulation (and the duration of different events that might occur), so that the effects of drift remain similar. For instance, with a rescaling factor of 10, the length of the simulation should be shortened by a factor of 10, as should the duration of events such as bottlenecks or expansions. Thus, rescaled simulations not only run faster per generation (because there are fewer individuals to process), but also run for a smaller number of generations. In the results section, we will show the effect of the rescaling factor on two summary statistics, along with the increase in the speed of the model. Because there are many complexities involved in rescaling, we recommend choosing this factor with great care, and cross-validating the results of downstream analyses by doing a small number of runs that are unscaled (or less rescaled, at least).

### 2.2 Simulation protocol

#### 2.2.1 Forward simulation

We now describe the protocol step by step. A schema in supplementary Figure S1 may help to understand the following section by giving an overview of the approach taken.

SLiM scripts can be called from the command line or run within the SLiMgui graphical modeling environment. Here we will define constant variables directly in the script, so that one can run the code in SLiMgui. When running the model at the command line, those constants could instead be passed to SLiM as `-d constant=value` command-line arguments; this is convenient to run a whole set of simulations with different parameters. In this example, we simulate 1000 generations of a population of 100 000 individuals, which have a chromosome of 2Mb, and a recombination rate of 10^-9^ bacterial recombination events per generation per base pair, with a mean recombination tract length of 10kb. The script begins with a block of code called an `initialize()` callback:

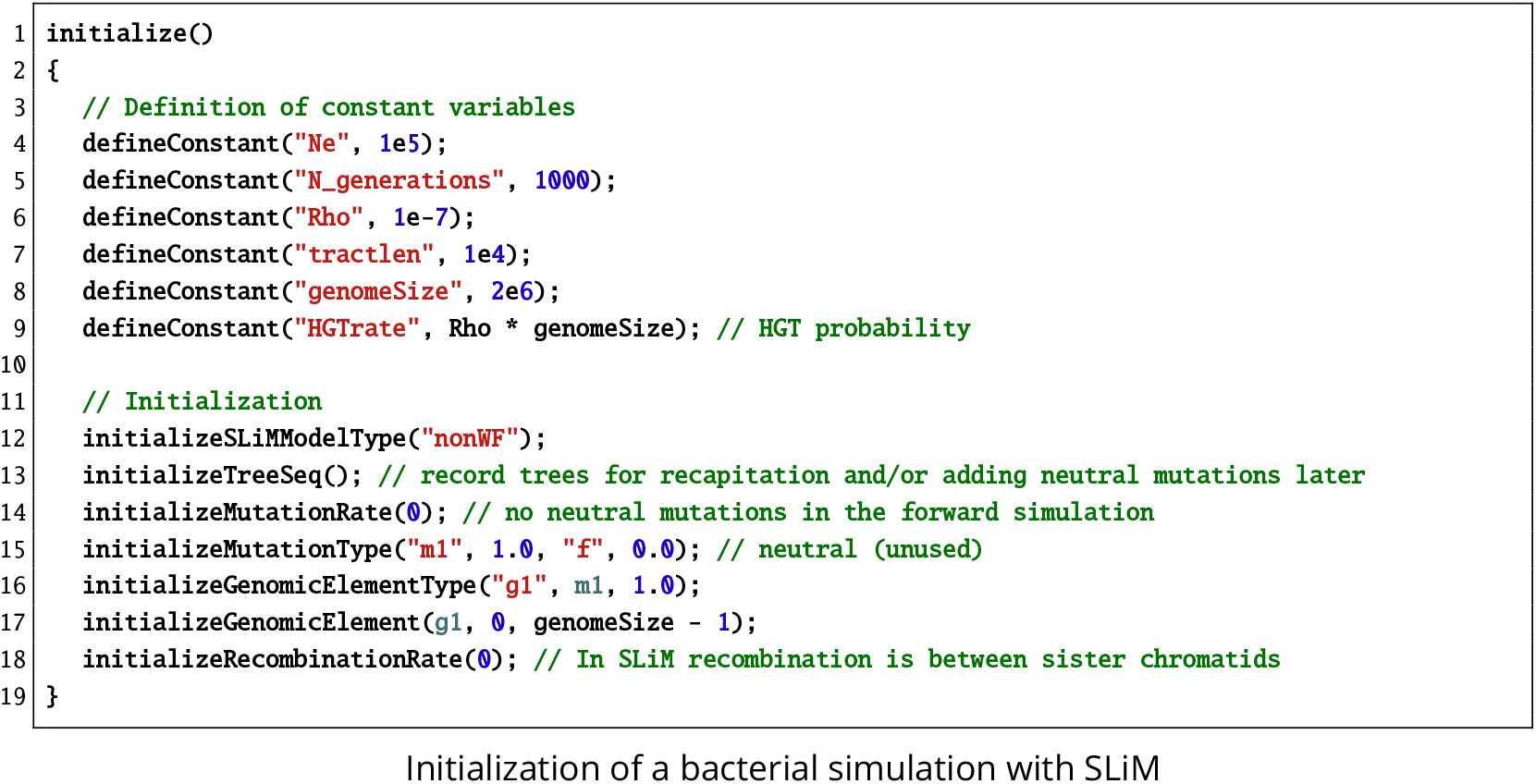
Initialization of a bacterial simulation with SLiM

Here we initialize the simulation using the nonWF model with tree-sequence recording, as explained in the SLiM manual. We set the mutation rate to zero because we will add neutral mutations laterwith `msprime`, after recapitation; we do notwantto forward-simulate neutral mutations, for efficiency. Importantly for bacteria, the (generic) recombination process implemented in SLiM should not happen, otherwise, because individuals in SLiM are diploids, our haploid bacterial chromosomes will recombine with the empty second chromosomes. Thus, the recombination rate should always be set to zero when simulating bacterial populations. Instead, we define a constant, `HGTrate`, that represents the probability of a given bacterium undergoing (homologous) HGT.

The population is created at the beginning of the first generation, as shown in the next snippet; other populations could be created here too:

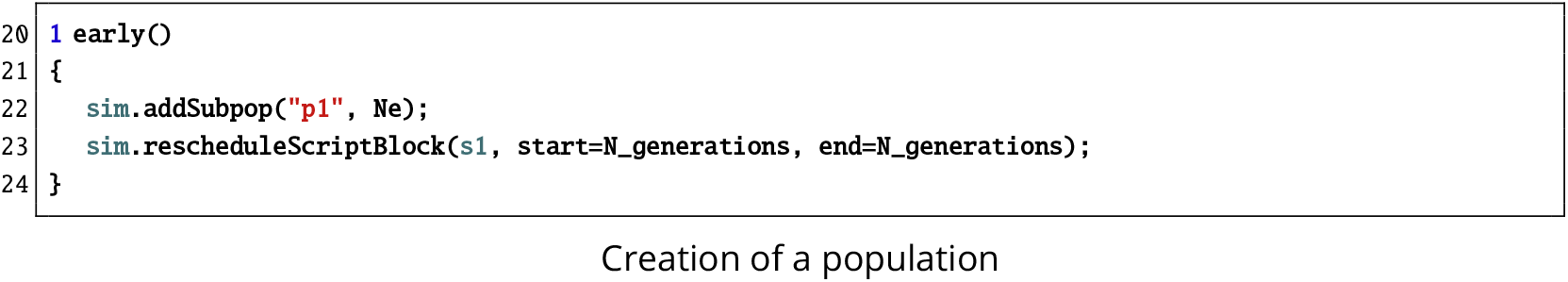
Creation of a population

In line 22, we add a subpopulation named p1 of size Ne. The next line is not specific to bacteria, but allows us to define the end of the simulation dynamically, governed by a parameter (`N_generations`). This is useful when comparing different rescaling factors, or when the endpoint of the simulation depends on other parameters or events.

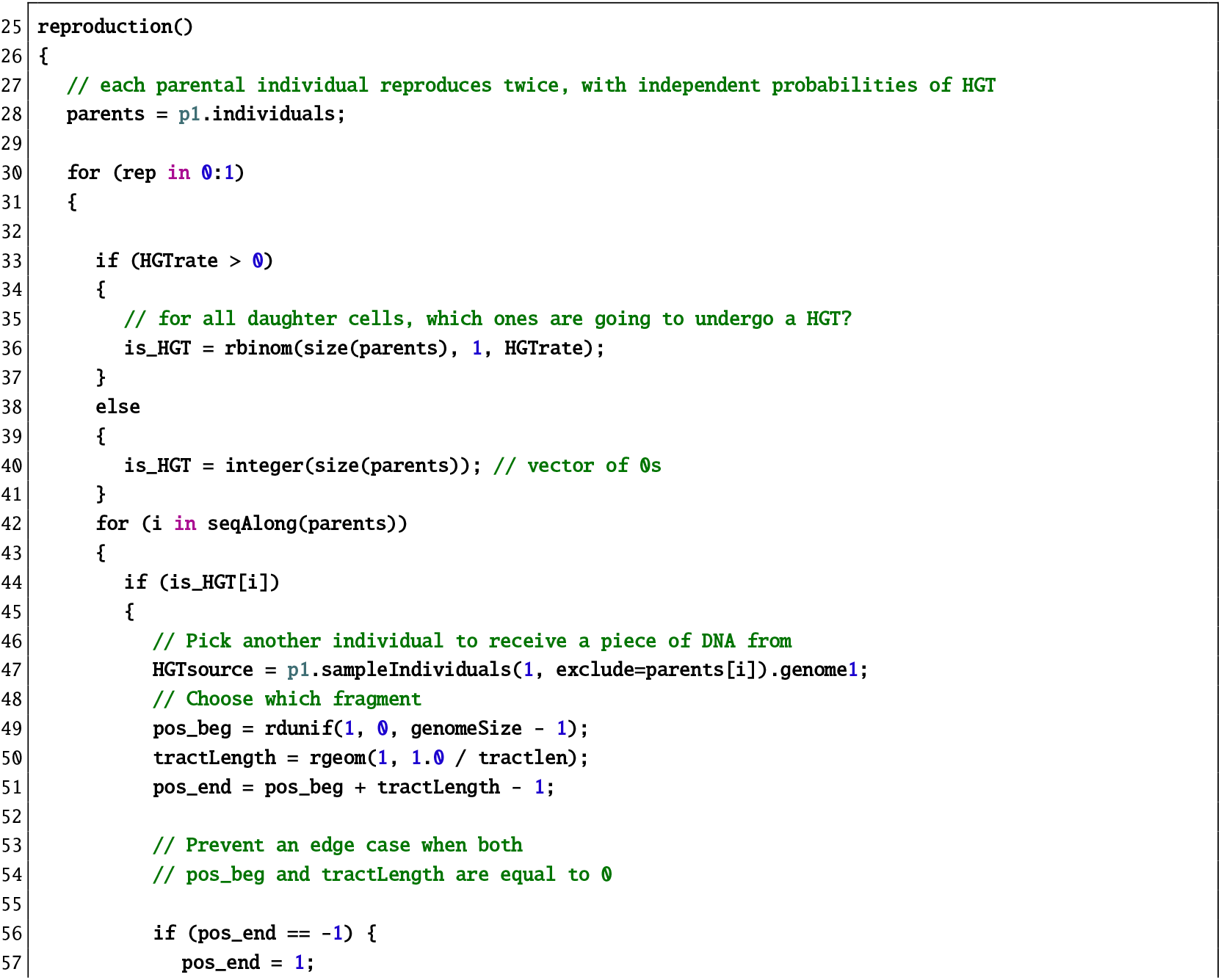
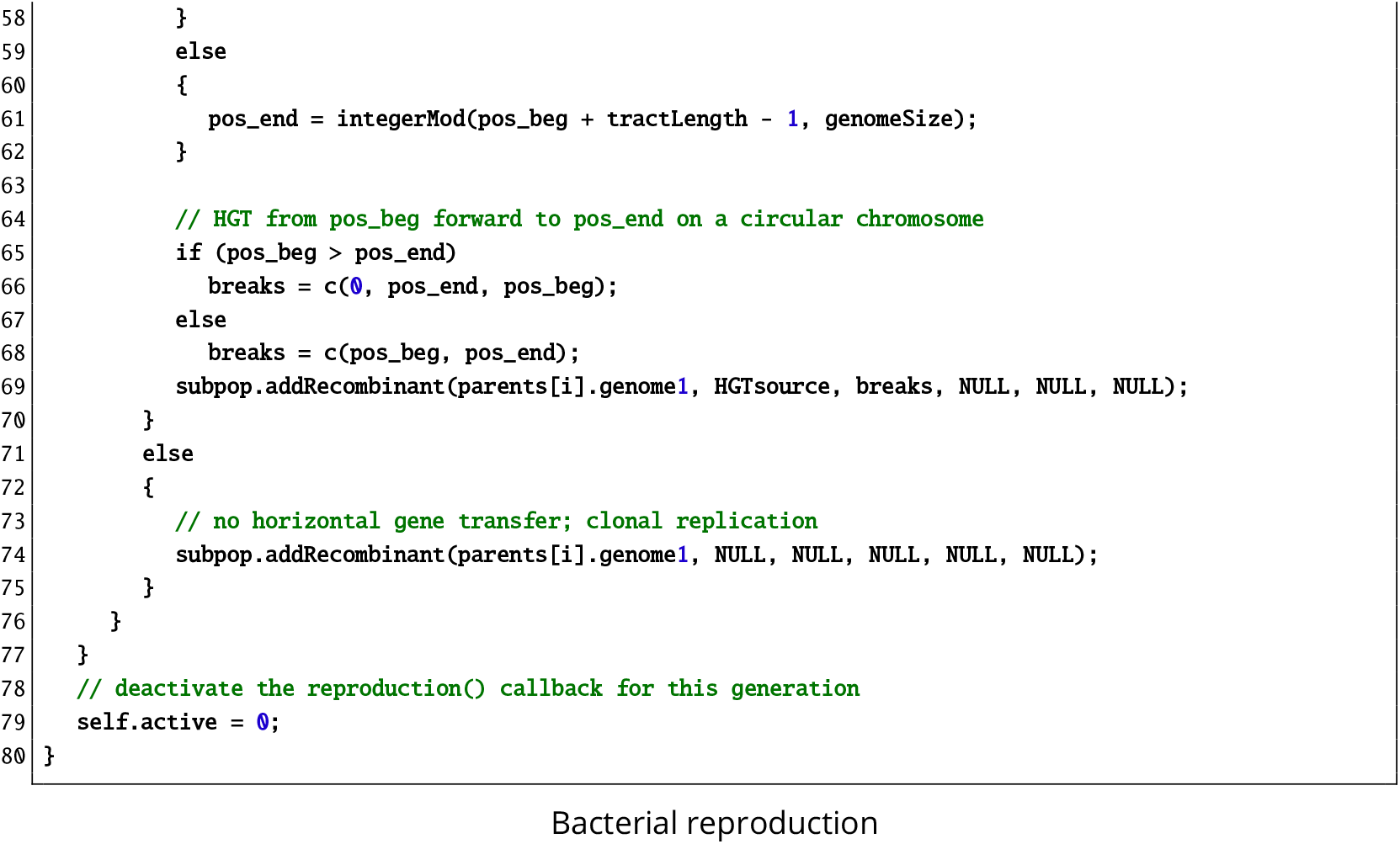
Bacterial reproduction

In each generation, SLiM calls `reproduction()` callbacks for each individual and the callback handles how that focal individual reproduces and generates offspring. Since we want to reproduce the whole population in one big bang (for efficiency, mostly), we override that default behavior by setting `self.active = 0`; at the end of the callback. As a result, this callback is called only once per generation and manages the reproduction of all individuals. We make each parent reproduce twice (`rep in 0:1`) to circumvent SLiM’s constraint that individuals cannot undergo a horizontal gene transfer event in the middle of their lifespan. By creating two clonal offspring, each can be part of a horizontal gene transfer event; had we implemented a single clonal reproduction, only one of the two daughter cells (the one that is not the parent) could have undergone HGT. Later in the script the parents are removed from the population (by setting their fitness to 0), such that in each generation, a bacterium reproduces, generating two daughter cells. We then decide which clones (see line 36 above) will undergo an HGT event by drawing from a binomial distribution, with the probability of HGT defined by the constant `HGTrate`, line 9. If an individual was chosen as a recipient for HGT, then the donor is picked randomly from the population (excluding the recipient); note that newly generated individuals are merged into the populations by SLiM at the end of reproduction, so a new daughter cell will never be an HGT source for another daughter cell. The DNA fragment that is going to be transferred is now defined by a starting position, drawn uniformly along the chromosome, and a length, whose value is drawn from a geometric distribution with mean equal to the tract length parameter (`tractlen`). Then, the `addRecombinant()` call creates a new daughter cell that is a clone of the parent, but with the recombination tract copied from the donor to the recipient. If the individual was not an HGT recipient, it is simply defined as a clone of its parent. Finally, as explained above, we deactivate this callback for the rest of the generation since it has just reproduced every parent.

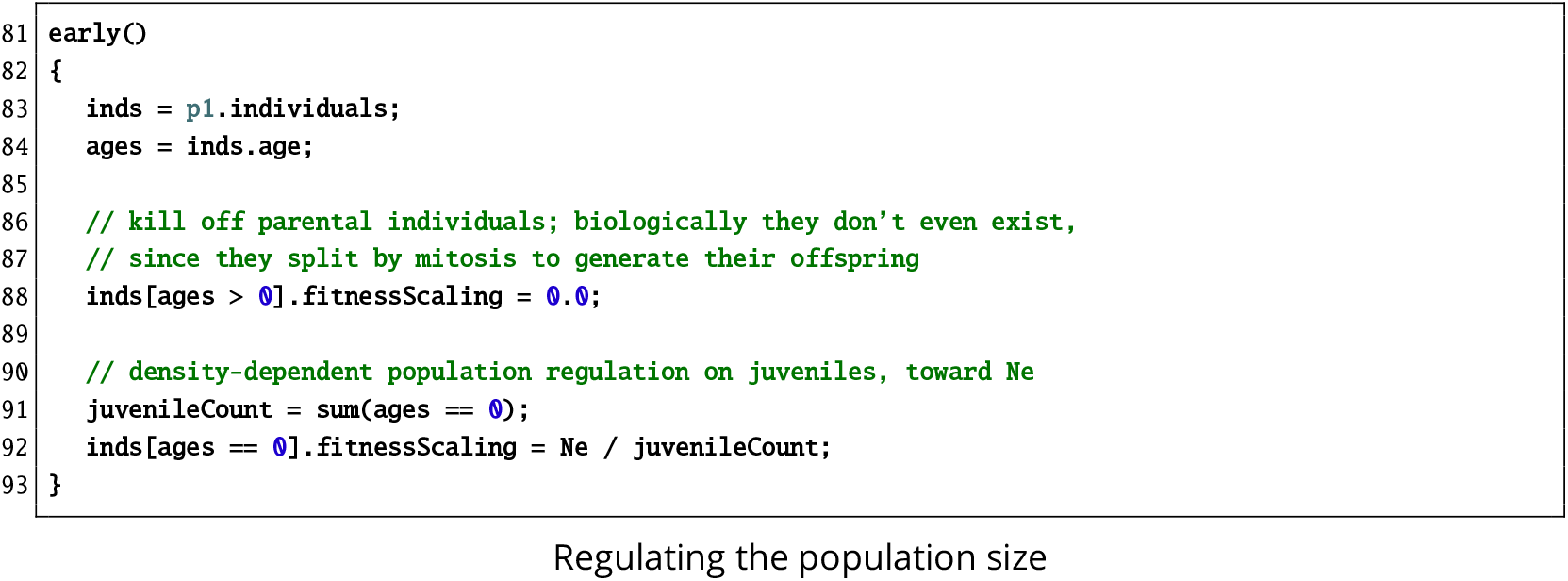
Regulating the population size

As we saw earlier, we had to clone each individual (parent) twice, to produce two new individuals (daughter cells/juveniles). To simulate mitotic cell division, we now remove the parents by setting their fitness to 0. In order to simulate a demographic scenario of constant population size, and because we are under the nonWF model where the size of the population is an emergent property (not a parameter as in WF models), we rescale the fitness of all juveniles so that the average number of individuals at each generation remains Ne. Before the next generation, SLiM will kill individuals based on their absolute fitness, which acts as a survival probability. Thus, at the start of the next generation we will have, on average, Ne individuals (with some stochastic fluctuation around that average).

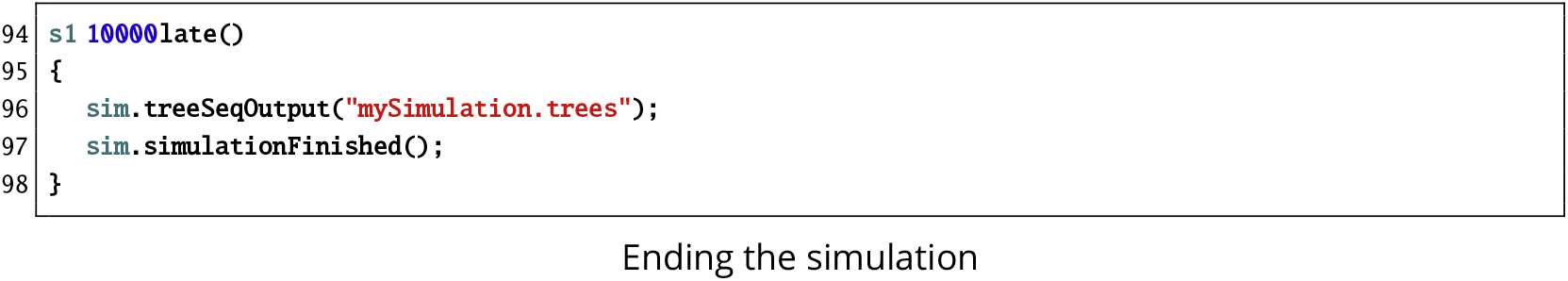
Ending the simulation

This script block, named `s1`, was rescheduled by `rescheduleScriptBlock()` in line 23, but a scheduled time for the block to execute – here 10 000 – has to be specified even though it will be overridden with `N_generations`. The value just needs to be high enough to avoid unintended execution of the block before it gets rescheduled; the time at which the unscaled simulation would end is typically a good choice, since it will never be too early. When the simulation is over, we output the tree sequence to a `.trees` file that we can work with in Python. In the next part we will show how to generate a burn-in period and genetic diversity `with msprime`.

#### 2.2.2 Recapitating and adding neutral mutations

When simulating with the nonWF framework, we efficiently obtain an initial population at mutation-drift equilibrium by performing a recapitation of the tree sequence, as explained earlier. So far, we have only forward-simulated the population while recording the tree sequence. Most likely, the simulation has not coalesced yet, because we did not run the simulation for at least 5Ne generations. We now recapitate the tree sequence, which runs backward in time, from the beginning of the forward simulation, to finish the coalescence process for our recorded tree sequence. Then, to obtain a matrix of neutral SNPs for the population at mutation-drift equilibrium – to compute summary statistics, for instance – the tree sequence can be manipulated in `msprime` with the help of `pyslim`, a Python interface between SLiM and `msprime`.

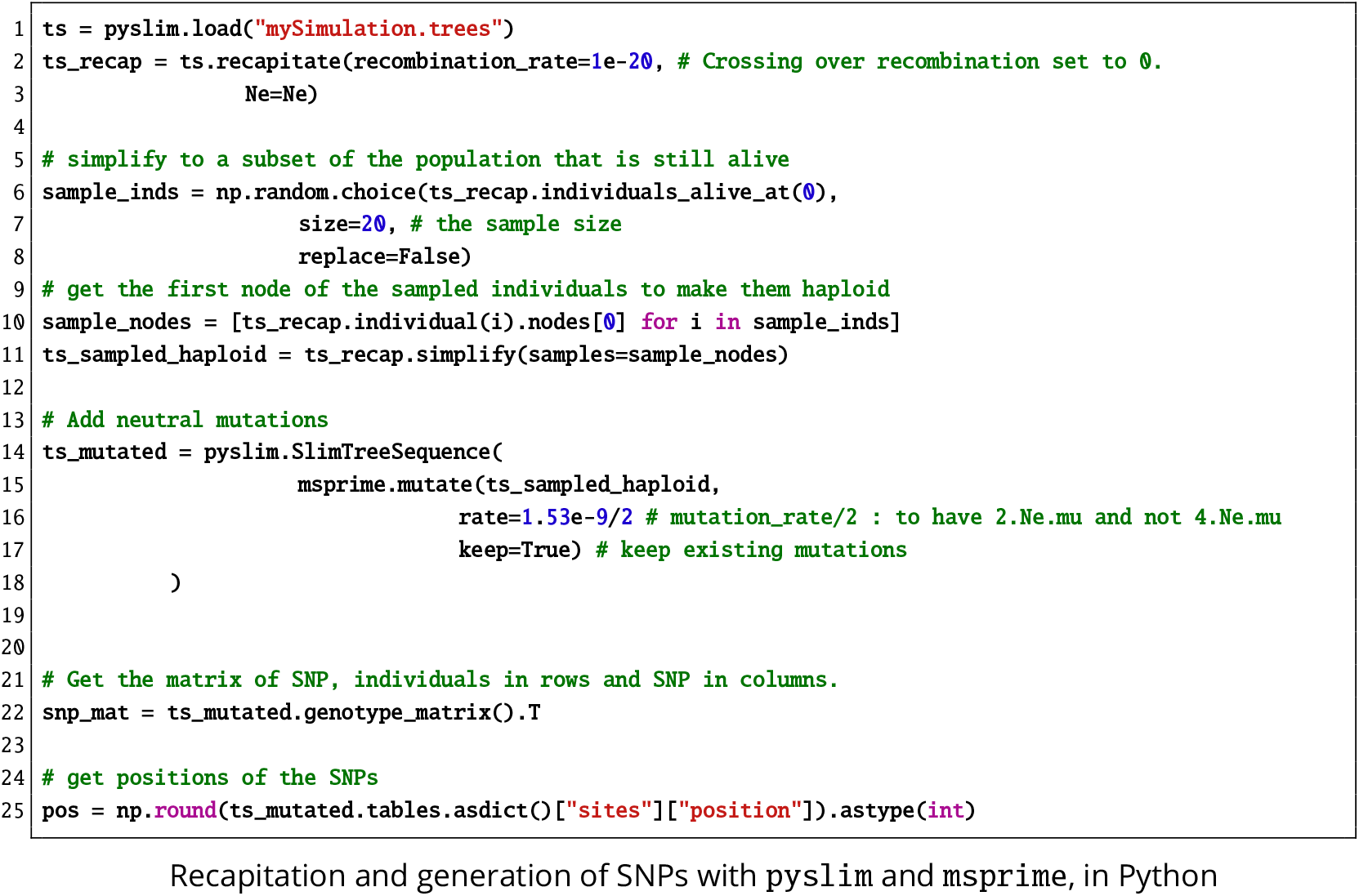
Recapitation and generation of SNPs with `pyslim` and `msprime`, in Python

First we load the tree sequence with `pyslim`, which returns a tree-sequence object. We then recapitate the branches that have not coalesced yet, with a very low recombination rate since `msprime` does not yet implement gene conversion. Since, for this protocol, we want to generate a matrix of SNPs for a sample of individuals, not for the whole population, we subsequently sample a random subset of extant individuals. We keep only the first node of each individual, corresponding to the first chromosome in SLiM where our haploid genetic material resides. Finally, we overlay neutral mutations on the resulting tree sequence. We have to divide the mutation rate by two to obtain the desired *θ* = 2 × *Ne × μ*, instead of *θ* = 4 × *Ne × μ* that `msprime` expects for diploids. At the end, we get a matrix of SNPs and a vector of corresponding positions, often used as input for inference methods [13], [16], [37]. This could easily be saved as an MS or VCF file if needed.

### 2.3 Simulations performed

To test the simulator, we ran simulations with parameters fitting the bacteria *Streptoccocus agalactiae* Clonal Complex 17, which is a major neonatal pathogen [38], [39]. We used a chromosome size of 2Mb, and we estimated the following parameters based upon data we found in the literature. The simulation spans 20,000 generations, which represents about 55 years of evolution for such bacteria in the wild, when using a generation time of 1 generation per day (as estimated for *E. coli* [40]). The mutation rate is set to 1.53 × 10^-9^ mutations per basepair per generation [38]. The recombination rate was set equal to the mutation rate, and the mean recombination tract length was estimated as 122 kb [41]. Note that the true recombination rate for *S. agalactiae* is probably lower [42], but in order to assess the correctness of the implementation of bacterial recombination, we chose to set it equal to the mutation rate and study the effects of varying it. The effective population size of this clonal complex was estimated to be around 140,000 individuals [38]. At the end of simulation, we sampled 20 individuals and built a matrix of SNPs, from which we computed summary statistics.

Simulations were run on a Dell R640 server rack with Intel Xeon Silver 4112 2.6GHz processors.

### 2.4 Simulating Bacteria on a Petri dish with antibiotic

To demonstrate the flexibility and scriptability of our SLiM bacterial model, we also present the results of a more complex and very different simulation scenario based upon our methods. It models bacteria growing on a Petri dish, seeded by 50 clones distributed randomly on the plate. Half of the plate contains an antibiotic that decreases a bacterium’s fitness, before density-dependent selection, from 1.0 to 0.47. However, a resistance allele can emerge through random mutation (ata rate of 10^-5^ per generation), and carrying this allele increases fitness back to 0.906 in the presence of antibiotic. In all cases there is a small cost for having the resistance allele, which leads to a reduced fitness of 0.98 for carriers of the resistance allele when antibiotic is not present.

Because this is a spatially explicit model the bacteria interact with their neighbors. In this type of model each bacterium has a given position in 2D space. Offspring appear near their parents, horizontal gene transfer occurs only between neighboring bacteria, and bacteria compete with their neighbors (which decreases the probability that a bacterium will divide under crowded conditions). More details can be found in the SLiM manual [35] in the section “Continuous-space models and interactions”. In this simulation with ongoing selection, neutral mutations are not tracked during forward simulation; they would be added during the recapitation phase. Only the resistance (beneficial) mutations are tracked here, since they are non-neutral and therefore influence the shape of the tree sequence.

At equilibrium, the probability that a given bacterium will divide is about 0.5, so half of the bacteria produce two offspring and die, while the other half produce no offspring and die, so the population size is then (stochastically) constant. The neutral mutation rate was set to 10^-9^, and the recombination rate was 10^-7^ with a mean recombination tract length of 500bp. The figures were generated using the random number seed 2049327378235, and snapshots were taken in SLiMgui. The code to reproduce this simulation can be found in the same repository: https://github.com/jeanrjc/BacterialSlimulations.

## 3 Results

We performed two sets of experiments to assess the performance and accuracy of our simulator. In the first experiment, we assessed the impact of rescaling the effective population size, *Ne*, in order to speed up the computation time. In the second experiment, we analysed the impact of varying the recombination rate and the mean recombination tract length, to better grasp their effects on the simulations. For both experiments, we monitored the running time and peak memory usage of SLiM, and assessed the quality of the simulations by comparing the site frequency spectrum (SFS) and the linkage disequilibrium (LD) with simulations obtained using `ms` [21] and `FastSimBac` [20], which are backward simulators implementing bacterial recombination (or gene conversion, in ms). For reference, we report runtime and memory footprint data for the backward simulators as well.

### 3.1 Impact of rescaling

We compared 9 different rescaling factors (RFs): 1 (no rescaling), 2, 3,4, 5,10, 25, 50, and 100. For RFs above 2, we generated 100 replicates for each RF and each SLiM model (WF, nonWF); for RF 1, 30 replicates were used, and for RF 2, 50 replicates. We generated 300 replicates when running `FastSimBac` and `ms` for comparison. Rescaling was not applied to backward simulators, since they do not simulate the entire population.

Without rescaling, the generation of a single replicate takes about a day (Figure 1). This is too long if one wants to run millions of simulations; however, it is possible to do a few such runs for other purposes, such as confirming that rescaling did not introduce a bias when implementing a new script. This might also be useful to test a method on a dataset produced without rescaling, since even minor artifacts introduced by rescaling could conceivably bias or confuse inference methods. When using a rescaling factor of 5, a simulation takes about 1 hour to run, which is practicable if one wants to run thousands of simulations on a cluster. With a factor of 25 or more, the running time is comparable to that of `FastSimBac` and `ms`, if not faster; `FastSimBac` is a bit slower than `ms`, probably because we used an additional `FastSimBac` script to create an `ms`-formatted output file. At 100 seconds or less per replicate, it is possible to generate about a million replicates in a few days or a week, on a typical computing cluster (using perhaps 100 cores). The time of the burn-in period is included here, and is not a limiting factor since it is faster than the forward-simulation period by about two orders of magnitude for rescaling factors smaller than 5 (supplementary Figure S2). The memory peak usage is fairly low (up to a few gigabytes without rescaling), allowing any modern laptop to run these simulations.

**Figure 1:**
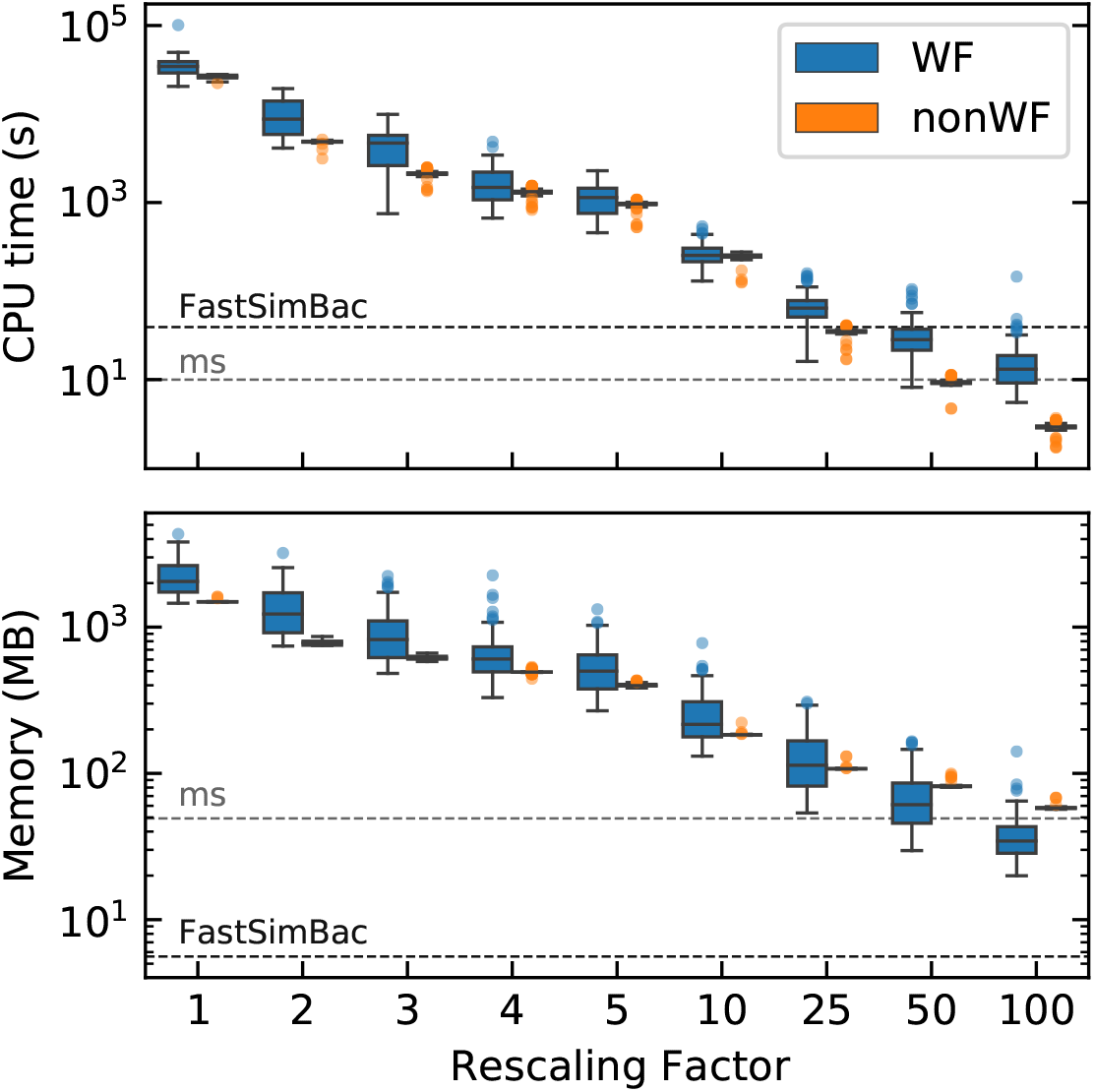
Distribution of CPU time and peak memory usage for different rescaling factors. For comparison with backward simulators, the dashed lines represent the average times for `FastSimBac (~54s)` and `ms (~14s)`. Rescaling does not apply to backward simulators. Note the log scale on the y axis; 10^5^ seconds is about 28 hours. Parameters used: chromosome size: 2Mb; *μ = ρ* = 1.53 × 10^-9^; *Ne* = 140k; 20000 generations. There are 30 replicates for RF 1, 50 replicates for RF2, 100 replicates for other RFs. Rescaling drastically reduces the computational time and memory usage, matching the performance of the coalescent simulators for sufficiently large RF.

Comparing WF and nonWF performance, we see that the nonWF model tends to be faster, especially at higher rescaling factors. This is due to the overhead of the burn-in step, which is slower in the WF models. Without rescaling, or at lower rescaling factors, the difference between WF and nonWF runtimes tends to disappear. Interestingly, the variance in time and in memory is lower for the nonWF version, which can help predict the resources needed for large runs. It is important to note that these performance metrics depend on the parameter values used (such as the recombination rate).

We then computed the normalized SFS produced by the different rescaling factors. The SFS represents the distribution of the frequency of derived alleles. Each bin (i) is given by 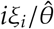, where *ξ_i_* is the count of SNPs having *i* derived alleles, and 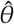 is an estimator of θ computed as the mean over the different bins (1/*n*∑*iξ_i_*). Because *iξ_i_* is an estimator of *θ*, the expected normalized SFS for a constant-size neutral population under the Wright-Fisher model is a flat line centered on 1 [43], [44].

Figure 2 shows the normalized SFS for 6 rescaling factors (see supplementary figure S3 for all RFs) with the expected standard deviation under the Wright-Fisher model without recombination [43]. `FastSimBac` and `ms` simulations are shown as a second control, in addition to the theoretical expectations (horizontal line at 1).

**Figure 2:**
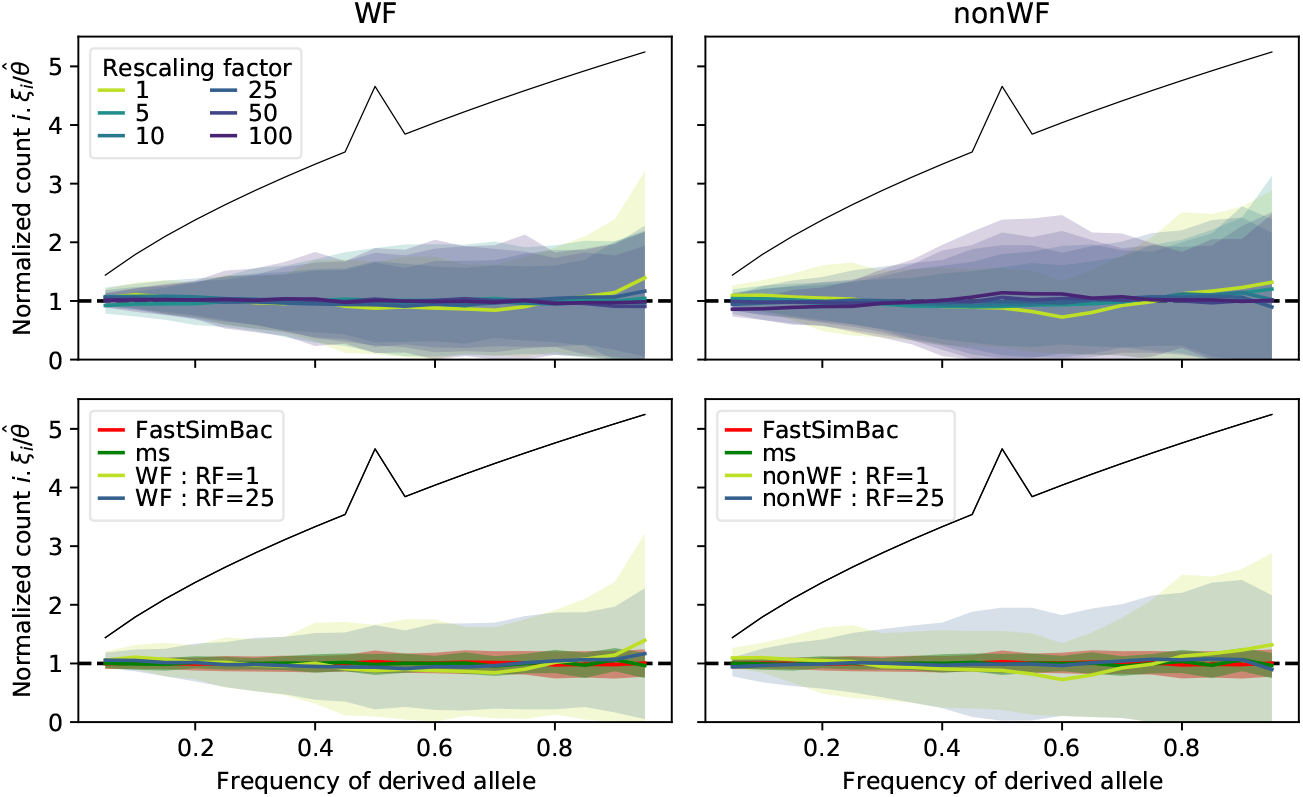
Normalized Site Frequency Spectrum (SFS) for four simulators and different rescaling factors. The left panels represent the SFS of the WF simulations under different rescaling factors (top left) and compared to the coalescent simulators (bottom left). The same information is shown on the right-hand side of the figure, butforthe nonWF simulations. The colored shaded areas represent one standard deviation (mean ± std). The horizontal dashed line at 1 indicates the expected average value and the black line the expected standard deviation, both under the WF model without recombination. Parameters used: chromosome size: 2Mb; *μ* = *ρ* = 1.53 × 10^-9^; *Ne* = 140k; 20000 generations. Rescaling does not affect the shape of the SFS and it matches that of the expected horizontal line at 1, is not different across rescaling factors, and is similar to the SFS obtained with the coalescent simulators.

We see that all experiments lead to the expected SFS, well within the expected standard deviation for linked loci. The smaller standard deviations, compared to the theoretical expectation, are not surprising since recombination is known to decrease the variance of the SFS [45]. Thus, rescaling factors up to 100 with this set of parameters do not affect the average SFS, which behaves correctly for WF and nonWF models.

Next, we assessed the impact of bacterial recombination on linkage disequilibrium (LD). The LD is measured by *r*^2^, which quantifies how much correlation (or linkage) there is between pairs of alleles separated by a given distance. We measured this correlation by subsampling pairs of SNPs, in 19 bins of increasing distances. The LD is represented as a function of the mean distance within each bin. We compared our results to the LD obtained with simulations from `FastSimBac` and `ms`. In figure 3 we observe that the LD for both WF and nonWF models is similar to that obtained with `ms` and `FastSimBac`, and does not seem to be affected by the rescaling factor. Unlike for the SFS, the expected LD and expected variation are hard to obtain and are beyond the scope of this paper. However, we know that the LD at very short distances should be close to the LD obtained in the absence of recombination. In figure 3, we used a shaded gray area to represent the range of LD without recombination at short distance. More precisely, it shows the mean +/− standard error of the mean for the four simulators without recombination. The full LD plot without recombination can be seen in supplementary figures S6 and S7. Note that in figure 3 a small difference between the backward and forward simulators can be seen, with the backward simulators tending to produce higher LD at short distances than the forward ones. This might be due to different implementations of recombination at short distances, or to a lack of recombination during the burn-in for forward simulation. Overall, however, we show that all types of simulations produced the expected LD at short distances and converged toward the expected *r*^2^ with free recombination of 1/*n* (dashed line) [46], [47].

**Figure 3:**
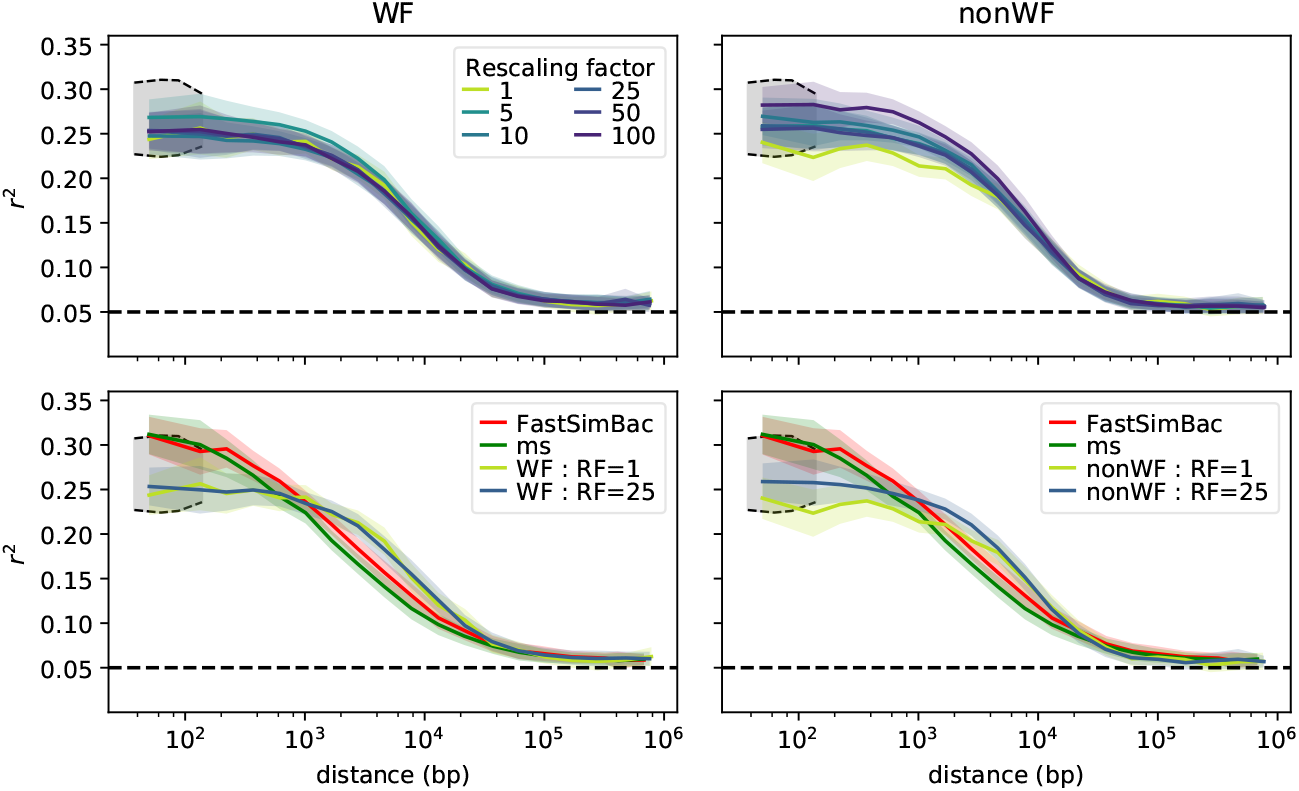
Linkage disequilibrium for WF (left) and nonWF (right) simulations under different rescaling factors (RFs) and for `ms` and `FastSimBac` (bottom). The horizontal dashed line indicates the expected *r*^2^ with free recombination when sampling 20 individuals (1/20). The colored shaded areas represent standard error of the mean, and the gray area toward the left of each panel represents the range of expected values at very short distances. Parameters used: chromosome size = 2Mb; *μ = ρ* = 1.53 × 10^-9^; *Ne* = 140k; 20000 generations. Rescaling does not affect the shape of LD (top), which matches that of coalescent simulators fairly well (bottom).

Overall, rescaling the simulations up to a factor of one hundred produces the expected SFS and LD, while allowing a drastic reduction in time and memory. This opens the possibility of running many simulations in a small amount of time, allowing the power and flexibility of forward simulation to be leveraged much more usefully in bacterial population genomics.

### 3.2 Impact of recombination

In this section we assess the impact of recombination with the same set of parameters used previously, with a rescaling factor of 25 across all of these runs. We compare simulations under three recombination rates (*ρ*|10, *ρ* and 10*ρ*, where *ρ* = 1.53 × 10^-9^) and three mean tract lengths (*λ*|100, *λ*|10, *λ*, where *λ* = 122 *kb*). These different recombination tract lengths cover the span of tract lengths found in bacteria where, depending on the mechanism of transfer, the size of the recombining region can range from approximately a 2kb fragment for transformation [48] to more than 100kb with conjugation [41]. It is worth stressing that for more realistic bacterial simulations, the mean tract length should represent the average for all recombination events, not only selected ones, otherwise the length might be overestimated [48]. We show here that a wide range of recombination tract lengths can be simulated. First looking at performance, increasing the recombination rate by a factor of 100 increases the runtime of the WF model 18-fold, but by only about 3-fold for the nonWF model (Figure 4 top). Higher recombination rates do require more memory, particularly when using the nonWF model, but with the rescaling factor used in this experiment it is still less than 1 GB (Figure 4 bottom).

**Figure 4:**
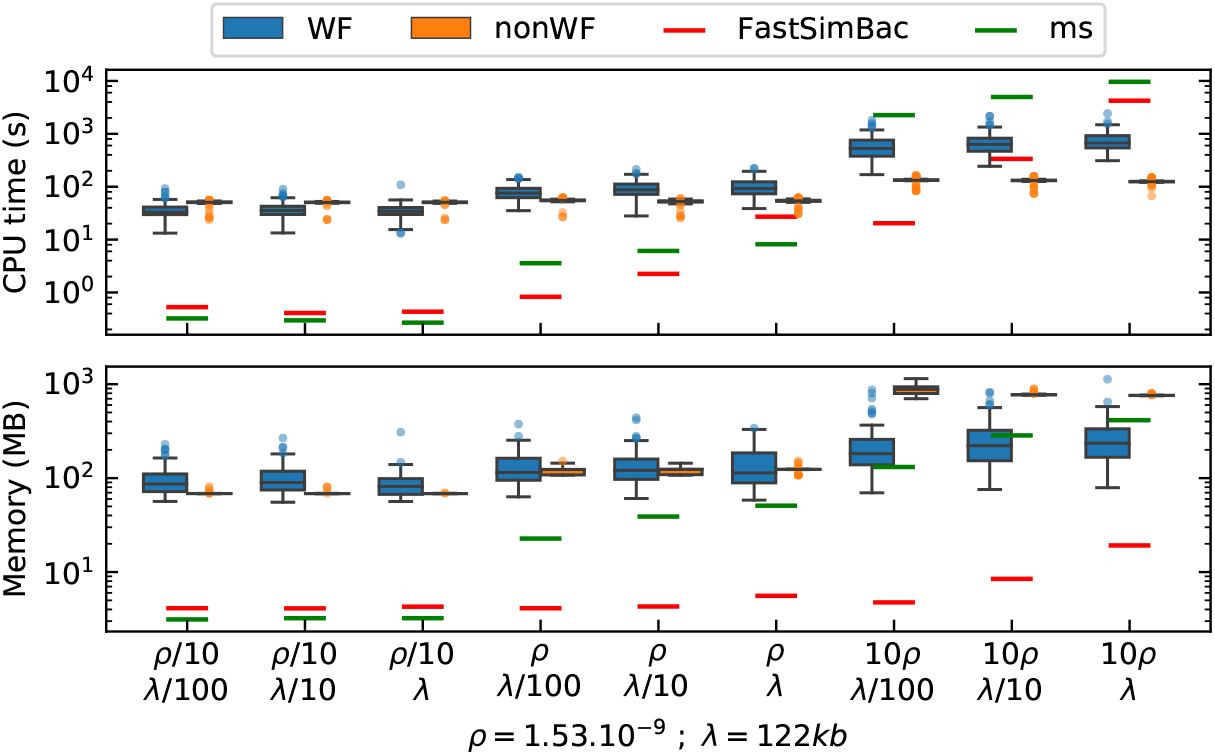
Distribution of the CPU time and peak memory usage for different recombination rates (p) and mean recombination tract lengths (*λ*). Parameters used: chromosome size: 2Mb; *μ* = 1.53 × 10^-9^; *Ne* = 140k; 30000 generations; RF=25. There are 100 replicates for each combination of *ρ* and *λ*. Computation time and memory usage increase with the recombination rate, but not with the recombination tract length.

The recombination rate thus has an important impact on the runtime of the WF simulations, but has much less impact on the nonWF simulations. The size of the recombination tract does not seem to significantly affect either runtime or memory usage. As expected, coalescent simulators are very fast at low recombination rates, but tend to struggle at higher recombination rates [20]. It takes them up to 10 thousand times longer to run when increasing the recombination rate by a factor of 100. Because of this, the simulations with `ms` and `FastSimBac` with 10*ρ* and *λ* were too slow, so we could only run 6 and 7 replicates, respectively, instead of a hundred. Other backward simulators may better handle higher recombination rates; however, ourfocus here is on the results of the simulations, not the efficiency of the simulators (a pointless comparison given the very different nature of coalescent simulators). The timing data for the backward simulators is just intended to give context for readers familiar with these software programs.

As in the previous experiment, we analysed the behaviour of our simulations with respect to the normalized SFS and the LD. In figure 5, we see that the SFS is distributed as expected (flat line centered at 1), independently of the simulator or type of simulation. Interestingly, we observed two expected theoretical results: the standard deviation of the simulated SFS at low recombination matches expectation [43], and the variance decreases as the recombination rate increases [45]. We see that for a given recombination rate, decreasing the recombination tract length has a similar effect as decreasing the recombination rate for a given tract length (moving between figure panels leftward is similar to moving between figure panels upward).

**Figure 5:**
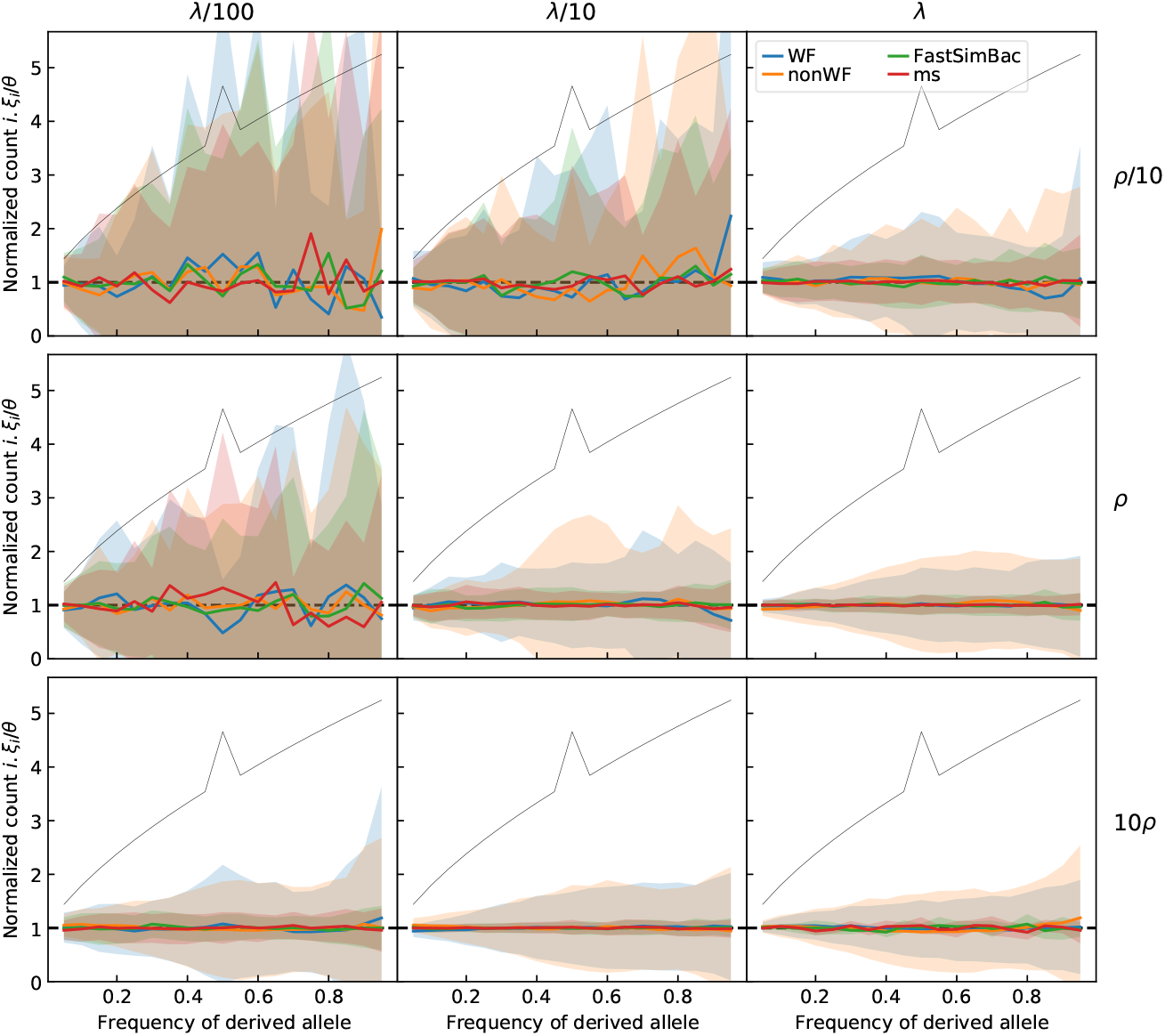
The normalized Site Frequency Spectrum (SFS) for different recombination rates (*ρ*) and tract lengths (*λ*). The colored shaded areas represent standard deviation. The horizontal line at 1 is the expected normalized SFS and the black line represents the expected standard deviation, both under the WF model without recombination. Parameters used: chromosome size = 2Mb; *μ = ρ* = 1.53 × 10^-9^; *λ* = 122kb; *Ne* = 140k; 30000 generations; RF=25. The observed SFS matches the expected horizontal line in all cases. Under low recombination their standard deviations also match the expectation, but the variance decreases with the recombination rate in accord with theoretical expectations.

In figure 6 the decay of LD with distance is similar when comparing all four types of simulations. We observe the same small discrepancy between coalescent and SLiM simulations as seen earlier, but only for a subset of the parameters. At low recombination rate, we recover the clonal frame, corresponding to the fact that bacterial recombination involves small patches of homologous DNA, rather than long stretches [49]. This means that positions on either side of an HGT patch will stay linked, and this explains the space between the line of the expected LD with free recombination and the LD curve at high distance, which is expected in bacteria [32]. A higher recombination rate or a longer recombination tract length tends to approximate the expected LD of an organism with recombination by crossing over rather than HGT.

**Figure 6:**
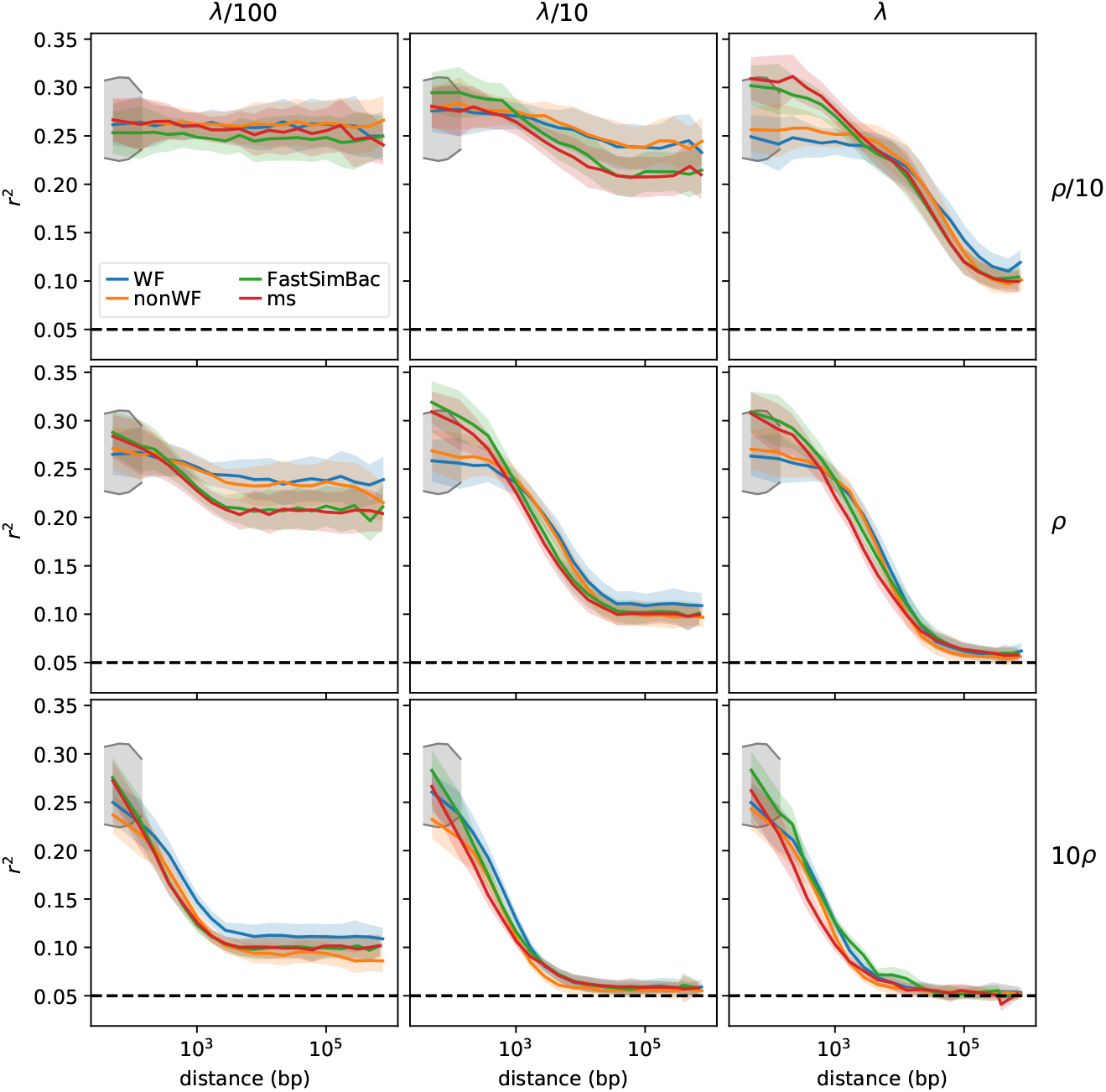
Linkage disequilibrium for WF and nonWF simulations with various recombination rates (*ρ*) and tract lengths (*λ*). The colored shaded areas represent standard error of the mean, and the gray area represents the range of expected values at very short distances. The horizontal black dashed line is the expected *r*^2^ with free recombination when sampling 20 individuals (1/20). Parameters used: chromosome size = 2Mb; *μ = ρ* = 1.53× 10^-9^; *λ* = 122kb; *Ne* = 140k; 30000 generations; RF=25. Various recombination rates and mean tract lengths produce similar patterns of LD between SLiM simulations and backward simulators.

Overall, changing the recombination rate and mean recombination tract length produced the expected statistical results. Even for the highest recombination rate, the runtime and memory requirements are still low enough to allow many simulations to be run (Figure 4), and if necessary, one might increase the rescaling factor (with proper validation and testing). Interestingly, with a high recombination rate the rescaled SLiM simulations were much faster than coalescent simulations. Finally, the nonWF model seems to have a more predictable runtime and memory footprint, which might be beneficial when computing resources are scarce.

### 3.3 Simulating bacterial growth on a Petri dish with antibiotic

In this paper, we mostly focus on results from a very simple population-genetic scenario. Here, however, we briefly showcase a radically different model, based upon our simple nonWF model, which might be of interest for evolutionary microbiologists. This model includes effects of explicit space on dispersal, competition, and genetic relatedness; this type of model may help with understanding the impact of environmental structure on a given evolutionary dynamic. For instance, a similar simulation framework was used to estimate the impact of a structured environment on resistance to phage and antibiotics [50]. In this toy scenario, we follow the growth of multiple colonies that are spread on a Petri dish as depicted in figure 7. The figure clearly shows how the antibiotic prevents the bacteria from spreading during a certain period and how the appearance of a resistance allele, despite being costly for its host in a neutral environment, eventually changes the spatial dynamics of colonization. This is obviously a basic model, and we are not interested in analyzing its results in any detail; instead, its purpose here is to show how easily more complex scenarios can be modeled in SLiM, based upon our simple bacterial protocol. Importantly, we still have access to the tree sequence of the population and we could thus recapitate the 50 individuals that started the simulations, overlay neutral mutations, etc., and perform further analysis.

**Figure 7:**
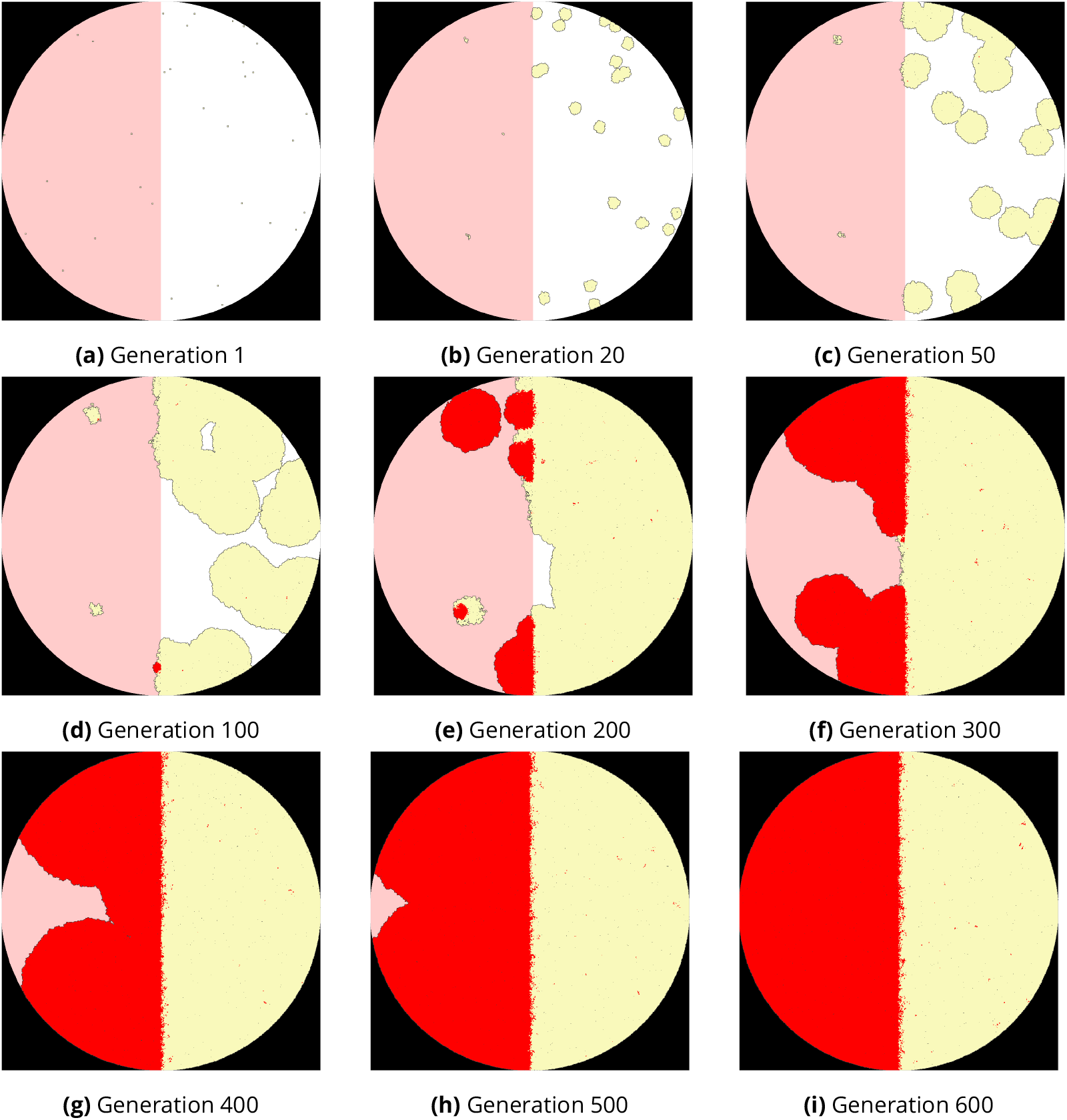
Representation of the simulated Petri dish at different generations. The pink area represents the half of the dish with antibiotic, while the white half has none. At the first generation, there are 50 colonies (single bacteria) spread randomly on the plate. As the simulation goes forward, we see those colonies growing, in yellow. They grow better in the right-hand half of the plate, due to the absence of antibiotic. Bacteria colored red have acquired antibiotic resistance. We see that the resistant bacteria later grow better in the left half of the dish. Because of the fitness cost of the antibiotic-resistance mutation in the absence of antibiotic, there are only a few red bacteria in the right half at the end of simulation despite their markedly higher mean fitness over the environment as a whole, showing the importance of spatiality for the outcome of the model. The code used to generate this simulation can be found at https://github.com/jeanrjc/BacterialSlimulations.

## 4 Discussion

We presented a step-by-step protocol for performing simulations in SLiM of a simple bacterial population, and one example of a more complex spatial model based upon our protocol. Although SLiM is not focused on bacteria, the simulations were shown to behave correctly, and ran in a reasonable amount of time. The basic models we presented were simple in order to draw attention to the particular techniques involved in simulating bacterial populations, but all of the model variation discussed in the SLiM manual – complex demography and population structure, selection, and so forth – can easily be added to this foundational model, as exemplified with our spatial model. Our simplified approach also allowed us to compare the accuracy of our implementation to theoretical expectations, and to other simulators for which substantially more complex scenarios would not have been possible. The final model, of bacteria on a Petri dish, certainly could not be run in any coalescent simulator.

These simulations were made both within SLiM’s Wright-Fisher framework and within the more individual-based nonWF framework, to showcase these two possibilities for the user. The recipes for our WF and nonWF models are freely available on our dedicated repository (https://github.com/jeanrjc/BacterialSlimulations), where we encourage everyone to propose their recipes for more complex scenarios. It might be worth mentioning why one would choose between SLiM’s WF and nonWF model types, since this fundamental choice will guide much of the model development that follows. The WF model is simpler in many ways: it involves more simplifying assumptions and less individual-level behavior. For example, population size in the WF model is automatically maintained at a set level, whereas the nonWF model requires you to write script that regulates the population size via mechanisms such as density-dependence or-appropriately for pathogens, perhaps – host mortality. Similarly, reproduction in the WF model is automatic, based upon fitness; high-fitness individuals reproduce more than low-fitness individuals, a fact that SLiM automatically enforces. In nonWF models, in contrast, fitness typically influences mortality, not fecundity, and reproduction is explicitly scripted to allow for greater individual-level variation in the modes and mechanisms of offspring generation. Writing a nonWF model is therefore a bit more complex and technical, and requires more details to be spelled out explicitly. Normally, nonWF models are a bit slower, but here the opposite was true; the slower implementation of the burn-in for the WF model, due to the incompatibility between tree-sequence recording and the WF implementation of bacterial recombination, meant that the WF model was slower. This is, in part, why we emphasized the nonWF model here; in this context, it really provides both greater power and flexibility, and better performance. However, the WF model remains simpler, conceptually and in its implementation; and if one wants fitness to affect fecundity rather than mortality it can be the more natural choice. These remarks are summarized in Table 1.

Currently, the only drawback of this simulator concerns the lack of recombination during the burn-in step. For the WF model, this is due to a technical limitation in *ms*; for the nonWF model, it is due to the current lack of gene conversion support in `msprime`. Implementation of gene conversion in `msprime` is in progress, and may be available soon. This will greatly improve the nonWF model, and will be trivial to add with a minor change to the recapitation step of the Python script. We will update our repository as soon as this feature is released in `msprime`. This lack of recombination during burn-in leads to a deficit in LD when forward simulation is too brief. In our runs, we see that after about 20 000 forward generations (about Ne/7 generations), LD and SFS match that of `ms` and `FastSimBac` (supplementary Figure S5 and Figure S6). If one wants to run a very short simulation (e.g. less than Ne/7) with burn-in, it might still be worth running at least Ne/7 generations more of forward burn-in in addition to a coalescent burn-in. The higher variance of the SFS observed in the experiments for the SLiM simulations, compared to `ms` and `FastSimBac`, might be explained by this lack of recombination during burn-in, since recombination decreases the SFS’s variance [45]. In short, for simulations that require burn-in (in order to start non-neutral dynamics at mutation-drift equilibrium, for example, or to obtain fully coalesced ancestry trees in Python), a few generations of neutral dynamics at the beginning of forward simulation are enough to recover the correct LD, at least in our simple model. (The necessary length of neutral forward simulation may be longer for other models, particularly with strong spatial structure.) Once gene conversion is added to `msprime`, this will not be needed any more.

We hope that our work here will stimulate a wave of development of simulation-based models for bacterial population genetics. We believe that this paper, combined with the hundred-plus models presented in SLiM’s extensive documentation, will allow anyone to create new scenarios for bacterial populations seamlessly. It is possible to simulate evolution in continuous space (such as in a Petri dish), to model nucleotides explicitly (including the use of FASTA and VCF files), to model selection based on external environmental factors such as the presence of antibiotics (and selection for resistance genes), and even to model within-host evolution using a single subpopulation for each host while modeling between-host transmission and infectivity dynamics; with the scriptability of SLiM almost anything is possible. We look forward to seeing the diverse research questions that the bacterial genomics community will explore with SLiM.

## Acknowledgements

We thank Peter Ralph, Eduardo Rocha, and Philippe Glaser for fruitful discussions. JC and FJ thank DIM One Health 2017 (number RPH17094JJP) and Human Frontier Science Project (number RGY0075/2019) for funding. The authors of this preprint declare that they have no financial conflict of interest with the content of this article. Guillaume Achaz is one of the PCI Evol Biol recommenders and is part of the managing board. Version 5 of this preprint has been peer-reviewed and recommended by Peer Community In Evolutionary Biology (https://doi.org/10.24072/pci.evolbiol.100123)

## Supplementary Figures

**Figure S1:**
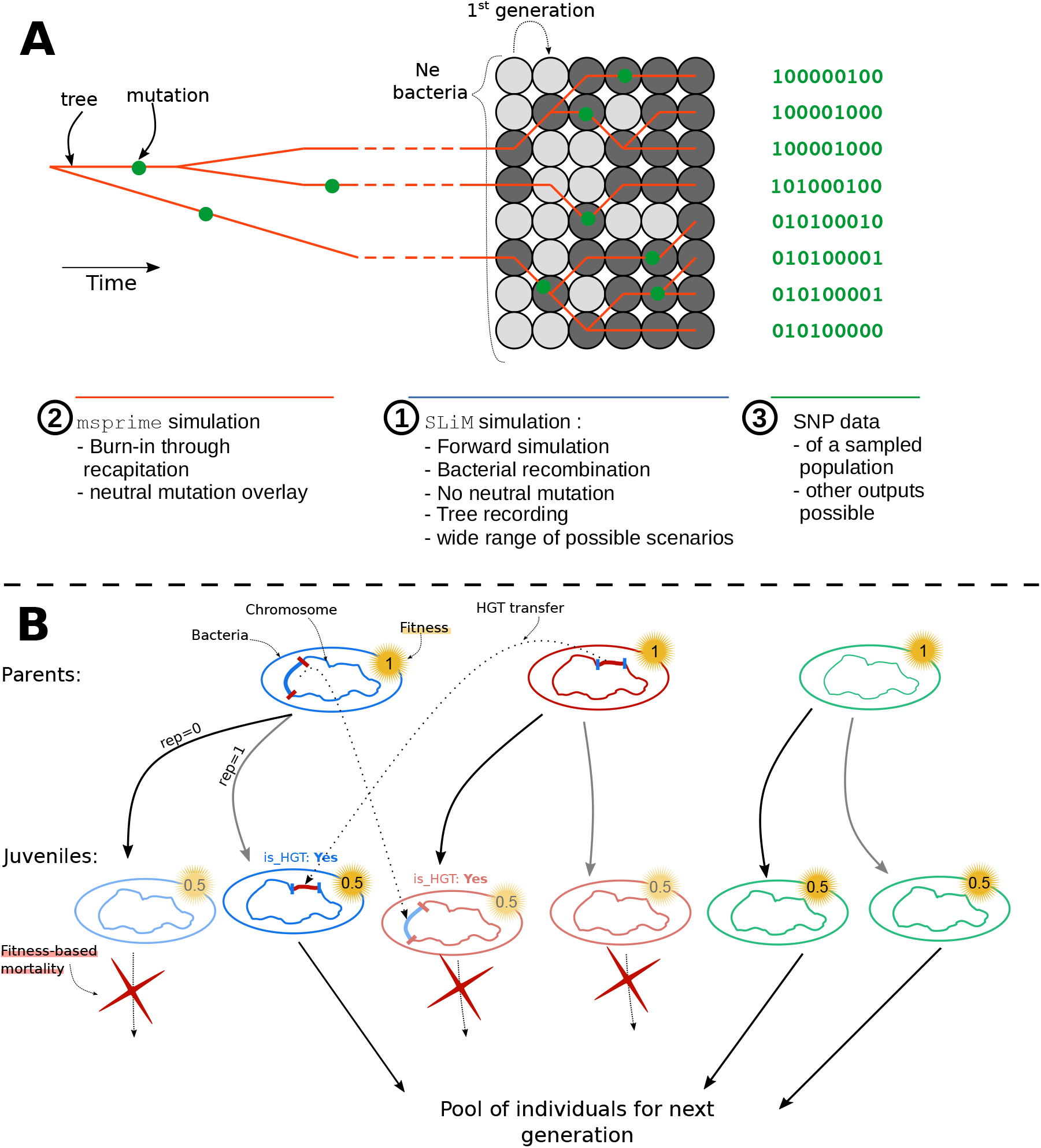
Schema of the nonWF simulation process described in this study. This schema is a visual companion to section 2.2 Simulation protocol, and is not self sufficient. (A) Step ①: forward simulation with SLiM of a population of bacteria with `Ne` individuals. The population at the time of sampling (on the right-hand side) will not have coalesced; here, for example, there are still three ancestors at the first generation (darker circles) and no single most recent common ancestor. Step ②: burn-in with `msprime` by recapitation of the tree sequence. The recapitation coalesces remaining branches back in time, acting as a burn-in period for the forward simulation. Then we can overlay neutral mutations on the fully coalesced tree sequence. Step ③: output SNP data for further analysis. Other types of data may be output, such as a VCF file, or a `.trees tree` sequence file. Panel (**B**) provides a simplified depiction of the reproduction process. Each parent produces two juveniles, and some of them will receive gene fragments (thicker chromosomal segments) from other parents by HGT. Parents that are going to receive HGT are drawn from a binomial distribution with the HGT rate as the probability (`HGTrate parameter`, L. 36 in the bacterial reproduction code snippet). The first position of the recombining fragment is drawn uniformly along the chromosome, and the second position is drawn from a geometric distribution with the mean tract length parameter (`tractlen parameter`, L. 50). The fitness is then adjusted by density-dependent selection, causing mortality (red crosses), such that on average the population size remains constant at equilibrium (see Regulating the population size’s code snippet, L. 81 – 92).

**Figure S2:**
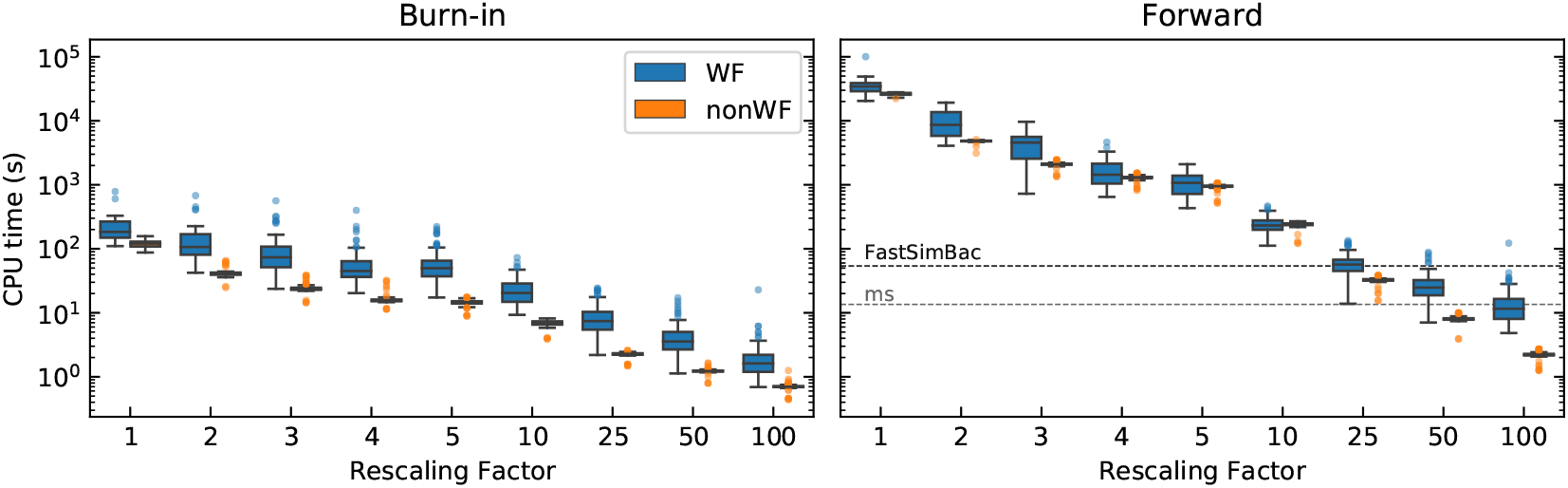
Computing time as in Figure 1, but split between the burn-in and forward simulation components of the total simulation time.

**Figure S3:**
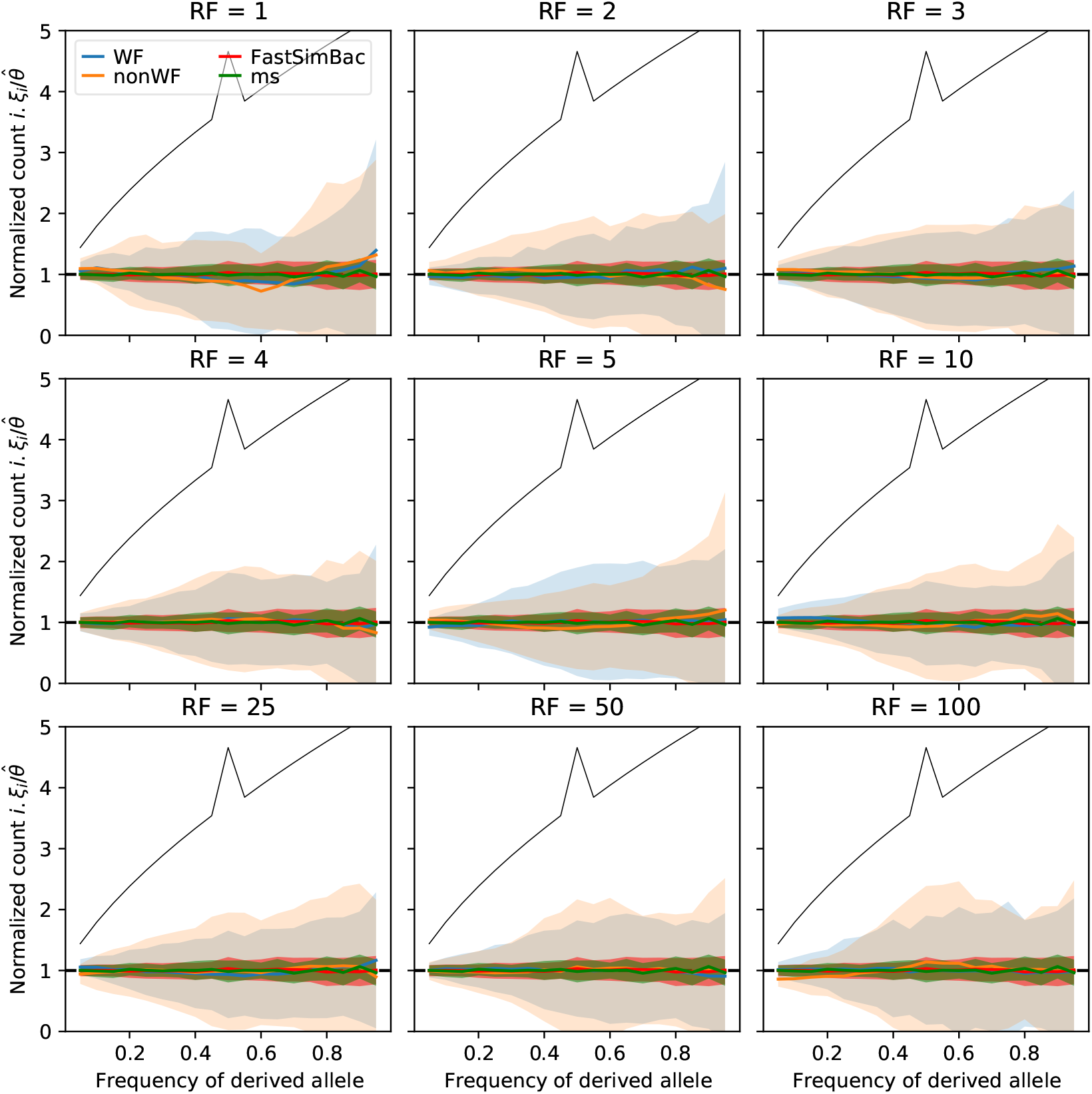
The normalized Site Frequency Spectrum (SFS) for different rescaling factors. The shaded area represents one standard deviation (mean ± std). The horizontal line at 1 is the expected normalized SFS and the black line represents the expected standard deviation, both under the WF model. Parameters used: chromosome size: 2Mb; *μ = ρ* = 1.53 × 10^-9^; *Ne* = 140k; 20000 generations. Rescaling does not affect the shape of the SFS and it matches that of the expected horizontal line at 1, is not different across rescaling factors, and is similar to the SFS obtained with the coalescent simulators.

**Figure S4:**
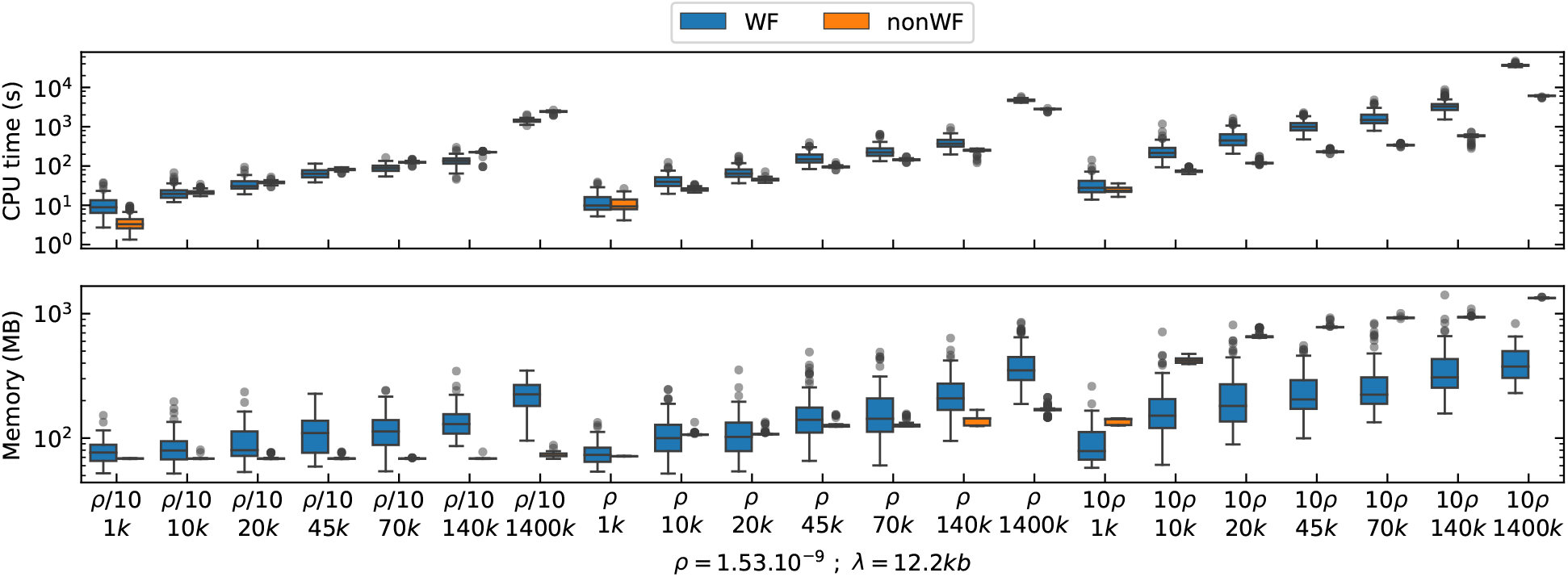
Computing time for different numbers of forward-simulated generations, with three different recombination rates. *Ne* =140 000, *λ* = 12200bp and RF=25.

**Figure S5:**
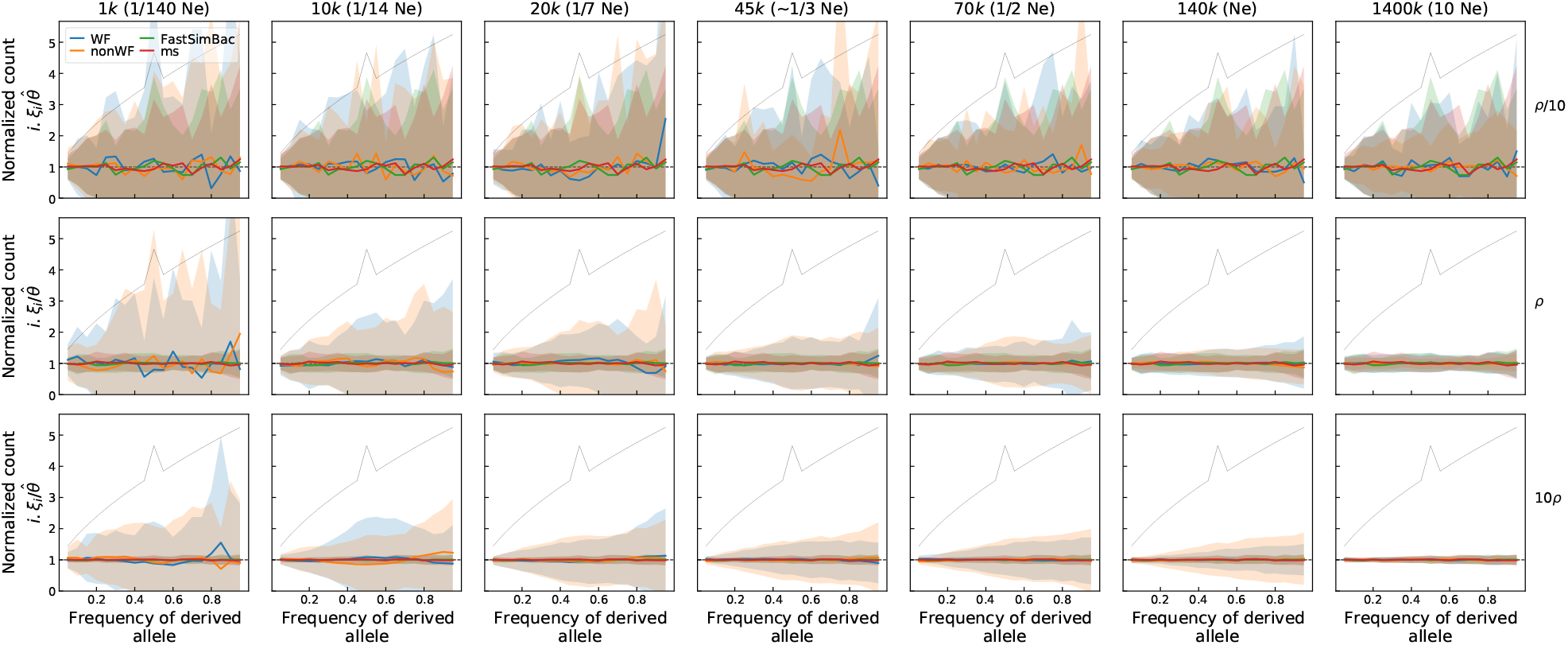
The normalized Site Frequency Spectrum (SFS) for different numbers of forward-simulated generations and recombination rates. The number of generations are indicated on the top of each column (e.g. 1k is 1,000 generations, and corresponds in unit of *Ne* to 1/140-th of *Ne*). The shaded areas represent standard deviation. The black lines represent the expected standard deviation for the Wright-Fisher model without recombination. *Ne* =140 000, *λ* = 12200bp and RF=25.

**Figure S6:**
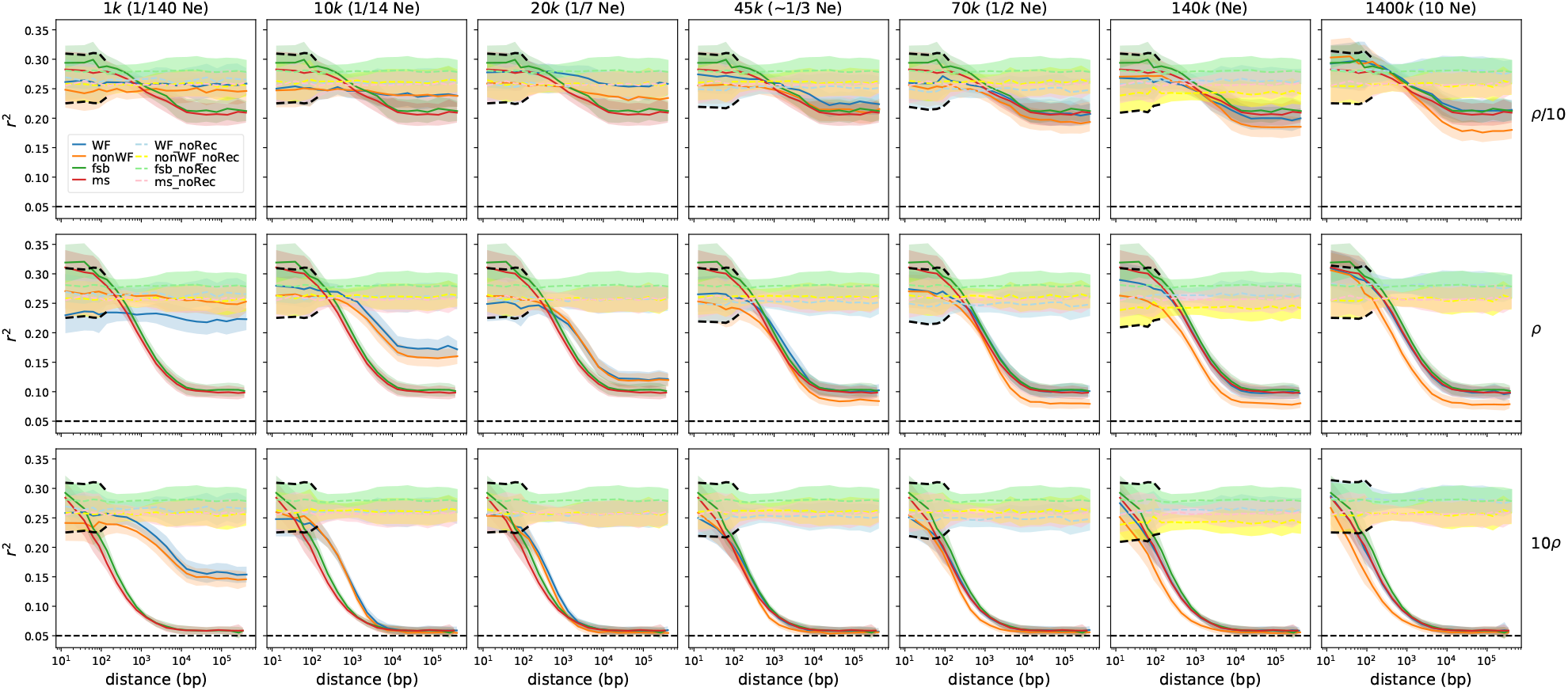
Linkage disequilibrium for WF and nonWF simulations with various numbers of forward-simulated generations and recombination rates (*ρ*). The number of generations are indicated on the top of each column (e.g. 1k is 1,000 generations, and corresponds in unit of *Ne* to 1/140-th of *Ne*). *Ne*=140 000, *λ* = 12200bp and RF=25. The horizontal dashed line is the expected *r*^2^ with free recombination when sampling 20 individuals (1/20). The shaded areas represent standard error of the mean; the standard deviation is 10 times larger (since we have 100 samples), as shown in figure S7 below.

**Figure S7:**
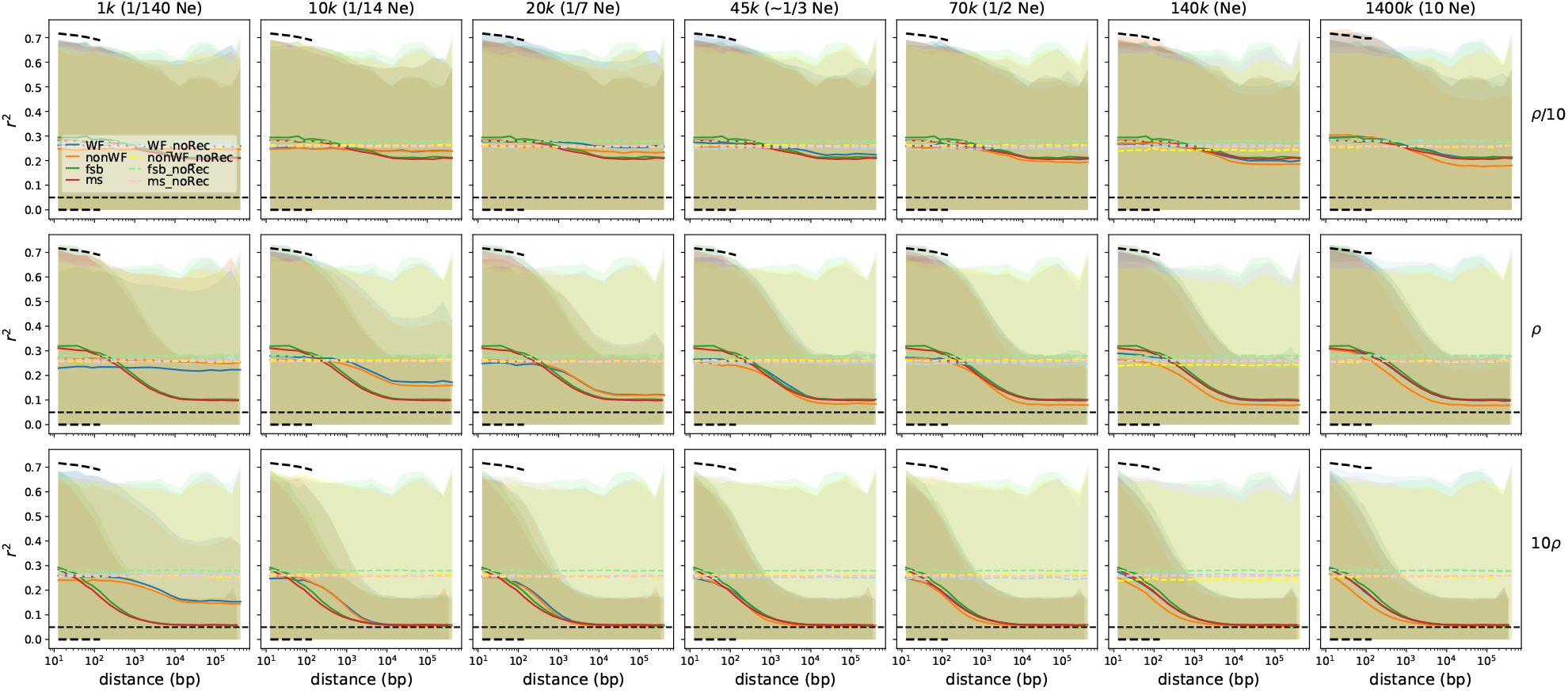
Same Figure as supplementary figure S6, but with shaded areas representing standard deviation instead of standard error of the mean.

## Annexes

### Demonstration of 5.N rule to reach mutation-drift equilibrium

Following Malecot’s derivation on heterozygosity [51], we have *H_t_*, the heterozygosity at time *t*, which can be expressed as follows:

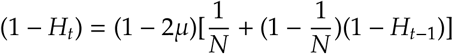

where basically, two individuals are identical at the previous generation (homozygosity, 1 – *H_t_*) if they coalesced (1/*N*) or if they were already identical and did not coalesce ((1 – 1/*N*)(1 – *H*_*t*-1_)). In both cases, no mutation should occur ((1 – *μ*)^2^ ~ (1 – 2*μ*)).

Rearranging the previous equation leads to:

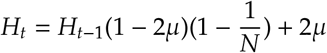

We can calculate the probability of heterozygosity at the equilibrium:

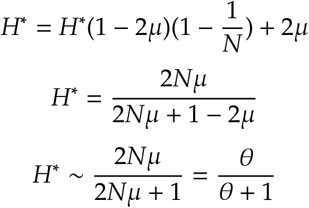

*H_t_* is an arithmetic-geometric sequence of the form *aH*_*t*-1_ + *b* that can thus be expressed as

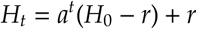

where 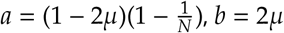 and 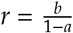.

Because |*a*| < 1, *H_t_* converges towards *r*; i.e., *r* is *H*\*, hence

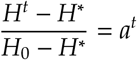

This ratio tends toward 0 as *H^t^* gets closer to the equilibrium. We want to estimate *t*_99_, the expected waiting time until the ratio falls down to 0.01, meaning that the heterozygosity is 99% closer to the equilibrium than when we started, i.e.,

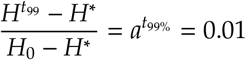

From this we get:

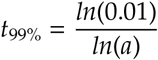

Because 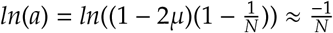 for small *μ* and large *N, t*_99%_ simplifies to

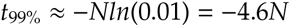

Thus after 5N generations the heterozygosity has almost reached its equilibrium, having progressed more than 99% of the way toward it, whatever the value of *H*_0_ (the initial heterozygosity).

